# Regional gene-environment association models improve insights into agents and targets of selection by climate in *Mimulus guttatus*

**DOI:** 10.64898/2026.09.28.755130

**Authors:** Ren N. Hamm, Michelle Stern, Ryan M. Boynton, Jack M. Colicchio, Victor Canta-Gallo, Jason P. Sexton, James H. Thorne, Nicholas J. Kooyers, Benjamin K. Blackman

**Affiliations:** Department of Plant and Microbial Biology, University of California, Berkeley, Berkeley, California, USA; Department of Environmental Science and Policy, University of California, Davis, Davis, California, USA; United States Geological Survey, California Water Science Survey, Sacramento, California, USA; Department of Life and Environmental Sciences, University of California, Merced, Merced, California, USA; School of Biological Sciences, University of Louisiana, Lafayette, Lafayette, Louisiana, USA; Department of Integrative Biology, University of California, Berkeley, Berkeley, California, USA

**Keywords:** genotype-environment association analysis, landscape genomics, genome-wide association analysis, climate adaptation, *Mimulus guttatus*, monkeyflowers

## Abstract

Genotype-environment association (GEA) approaches identify genetic variants that likely contribute to adaptation along environmental gradients. These associations can inform predictive models of maladaptation risk and evolutionary rescue in future climates. However, because population structure and climate parameters often co-vary, tradeoffs arise between accounting for population structure to reduce false positive associations and removing true positives. Here, we examine whether conducting GEA analyses at multiple spatial scales provides more informative associations by mitigating confounds between population structure and environment, and we evaluate what factors influence how congruent GEA results are across regions. We performed GEA analyses for climate parameters on genome resequencing data obtained for 110 annual populations of the common monkeyflower, *Mimulus guttatus*. We find that more genetic variation was specifically attributable to environmental variation in regional GEA models compared to a full-range model, and that agents of selection differ considerably in importance among regions and range-wide. Few genetic targets of selection shared across regions were detected, yet all regions possess ample standing genetic variation for most environment-associated variants, suggesting that regional differences in climate breadth may be a primary cause of this incongruence. In combination with a genome-wide association study for several climate-associated phenotypes, we also find that regulators of flowering repressors in the *FLC/MAF* gene family often harbor adaptive variation in this species. Together, our results suggest that GEA analyses of widespread species considerably benefit from examining both range-wide and regional trends to more fully capture and understand the agents and targets of climate adaptation.

## Introduction

A century of studies provides empirical evidence that local adaptation is relatively common in widespread species (Hereford, 2009; Leimu & Fischer, 2008), especially across environmentally heterogenous regions such as the western United States (Baughman et al., 2019; Clausen et al., 1940). However, it remains challenging to connect specific agents of selection (i.e., the selective pressures) to the particular targets of selection (i.e., the phenotypes and genetic variants) that afford local genotypes their fitness advantages compared to genotypes from elsewhere (Bomblies & Peichel, 2022; DeMarche et al., 2022). Such mechanistic knowledge is necessary to predict future responses to selection, and thus critical information for preserving natural populations when mounting evidence indicates the pace and severity of climate change is outstripping the capacity of many plant species to adapt effectively (Kooyers et al., 2025). Indeed, adaptational lags, where native populations have lower fitness than geographically-distant populations characterized by historical climates more akin to contemporary local conditions, have now been identified across seed plants (Anderson & Wadgymar, 2020; Browne et al., 2019; Fréjaville et al., 2020; Kooyers et al., 2019; McGraw et al., 2015; Pennington & Sexton, 2026; Quarles-Chidyagwai et al., 2026; Wilczek et al., 2014).

Moreover, many surveyed species demonstrate below replacement population growth rates in at least a portion of their species range (Anderson et al., 2025; Campbell, 2019; Inouye, 2008; Krushelnycky et al., 2013; Reed et al., 2021; Sheth & Angert, 2018; Zettlemoyer et al., 2024). Understanding the alleles and traits that have historically conferred fitness advantages across climatic gradients can help predict which species may be able to persist by evolving apace with changing climate, either unaided or with targeted management interventions (Aitken et al., 2024; Anstett et al., 2026; Breithaupt et al., 2026; Capblancq et al., 2020).

Genotype-Environment Association (GEA) analysis is a landscape genomics approach that has become an increasingly common means for exploring how spatially varying selection imposed by climate factors has historically shaped clinal patterns of genetic variation (Carvalho et al., 2021; Gibson & Moyle, 2020; Grabowski et al., 2025; Hancock et al., 2011; Hecht et al., 2015; Lasky et al., 2012). Multivariate methods like Redundancy Analysis (RDA) detect environment-associated genetic differentiation by modeling multiple environmental factors simultaneously (Capblancq & Forester, 2021; Fitzpatrick & Keller, 2015; Forester et al., 2016, 2018), but connecting these composite signals back to specific environmental agents of selection may not always be straightforward. In contrast, univariate methods like Latent Factor Mixed Models (LFMM) test associations between individual environmental variables and allele frequencies (de Villemereuil et al., 2014; Frichot et al., 2015). However, significant results come with the caveat that the actual causal environmental factor responsible for the association may be another unexamined but correlated variable. Nonetheless, by applying both of these GEA approaches and identifying overlapping hits, it can be possible to enrich for true signals of local adaptation (Bekkevold et al., 2019; Carvalho et al., 2021). Complementing GEA with genome-wide association (GWA) studies to connect genotype to phenotype for known locally adaptive traits may further contextualize why certain alleles are favored under particular environmental regimes (Lasky et al., 2023; Lotterhos, 2023).

False positive signals in GEA analysis are often controlled for by including population structure and/or geography as covariates (Hoban et al., 2016; Rellstab et al., 2015). However, because patterns of population structure and geography often covary with environmental factors, doing so often comes at the cost of eliminating true positives as well. Some studies therefore forego controlling for population structure and instead enrich for true positives by looking for congruent signals across different regions or taxa (Capblancq et al., 2023; Gutiérrez-Guerrero et al., 2024; Yeaman et al., 2016). A key drawback of this approach is that many true positives may be ignored, as there are diverse biological reasons why congruence may not be expected across regions. For instance, the range or selective importance of individual or composite environmental variables may not be equivalent across regions, allelic variants common in one region may be absent or rare in other regions, and/or different alleles at the same or different loci may have substitutable effects along similar environmental gradients (Lasky et al., 2023; Lotterhos, 2023; Rellstab et al., 2015). In addition, differences in statistical power due to variation in sampling scheme or due to differences among regions in how population structure covaries with the environment may result in incongruence (Coop et al., 2010; De Mita et al., 2013; Lotterhos & Whitlock, 2015). Thus, absence of congruent GEA results across samples or studies may often be expected and warrants attention as an opportunity to better understand the genetics and ecology of local adaptation. Moreover, incorporating region-specific GEA hits within predictive models developed for species conservation, e.g., use of genetic offsets, could aid in developing more targeted and successful management practices (Aitken et al., 2024; Capblancq et al., 2020).

Here, we leverage natural populations of the common yellow monkeyflower, *Mimulus guttatus* [syn. *Erythranthe guttata* (Lowry et al., 2019; Nesom et al., 2019)], as an ecological model to identify the agents and targets of selection by climatic factors across multiple geographic scales. *Mimulus guttatus* has a large native range, spanning Western North America from the Aleutian Islands to Mexico, and occupies diverse habitats (Grant, 1924; Vickery Jr, 1978; Wu et al., 2007; Yuan, 2019). Within California and Oregon, populations can be found over broad elevation and latitude gradients corresponding to a wide array of ecoregions (Griffith et al., 2016; Kooyers et al., 2015; Twyford & Friedman, 2015). Across this heterogenous landscape, *M. guttatus* exhibits both continuous and regionally discrete patterns of population structure and has several phenotypically distinct ecotypes (DeMarche et al., 2016; Ivey et al., 2023; Twyford et al., 2020). Coastal perennial and inland perennial populations benefit from persistent water availability, with coastal populations also adapted to endure substantial salt stress (Lowry et al., 2008; Plunkert et al., 2026; Toll et al., 2026). Inland annual populations inhabit seeps, drainages, and other habitats where ephemeral water availability leads to growing seasons that end in a terminal drought, and while the soils of some populations are fed by snowmelt, other populations rely solely on seasonal rainfall for moisture (FitzPatrick et al., 2023; Hall & Willis, 2006; Mojica et al., 2012).

Local adaptation to climate is widespread across annual *M. guttatus* populations. Heritable variation for many morphological and phenological traits is distributed along clines in temperature and precipitation and often related to growing season timing, and different populations have evolved toward different drought resistance strategies in response to historic and contemporary conditions (FitzPatrick et al., 2023; Kooyers et al., 2015, 2021). Yet despite having characteristics favorable for rapid evolution, including annual reproduction, high fecundity, and very high levels of standing genetic variation (Lovell et al., 2025; Puzey et al., 2017; Wu et al., 2007), common garden experiments indicate this species may still be experiencing an adaptational lag to climate change (Kooyers et al., 2019) with some populations suffering complete mortality during an extreme climate event (McDonald et al., 2023).

Therefore, by identifying genomic targets of selection and associating them with climatic agents of selection or variation in locally adaptive traits, we can begin to understand what genetic variants could be valuable for promoting population persistence in response to climate change. In addition, the extensive environmental heterogeneity and population structure of these populations make this species a potentially instructive system for testing expectations and explanations for (in)congruence in GEA results.

A previous small-scale GEA study that we conducted for a transect in Oregon highlighted candidate genes bearing several signatures consistent with adaptation to locally important agents of climatic selection (Colicchio et al., 2021). Therefore, we substantially expanded this whole genome resequencing dataset to include annual *M. guttatus* individuals sampled from 110 native populations throughout Oregon and California that span the breadth of climatic diversity within that range [Figure 1]. By performing multivariate and univariate GEA analyses as well as a GWA study with these data [Figure 1a], we sought to answer the following questions: (1) how do geographic, demographic, and environment structure genetic variation at range-wide and regional spatial scales?, (2) what are the primary *agents* of selection, (3) how congruent are the *targets* of selection observed across regions, and what biological or methodological factors may explain incongruence?, and (4) what do the sets of environment-and phenotype-associated genetic variants reveal about important *targets* of selection across the range?

**Figure 1:**
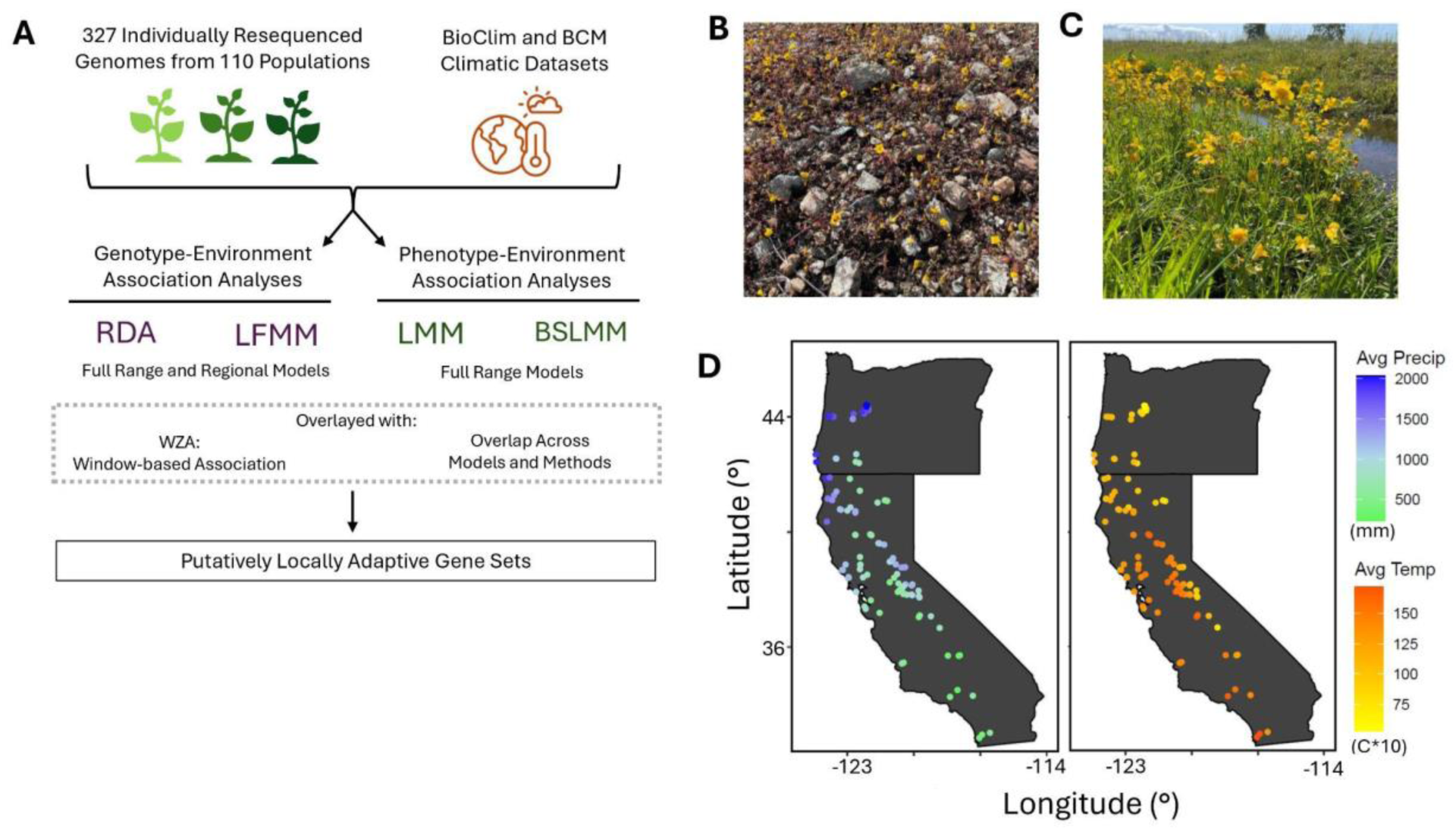
Experimental Plan and Sampling Scheme. (A) Pipeline for Genotype-Environment Association (GEA) and Genome-Wide Association (GWA) analyses to identify putatively adaptive alleles. GEA methods include Redundancy Analysis (RDA) and Latent Factor Mixed Models (LFMM); GWA methods include Linear Mixed Models (LMM), Bayesian Sparse Linear Mixed Models (BSLMM). (B-C) Images of annual *Mimulus guttatus* observed in (B) the Red Hills, Jamestown, CA and Galt, CA (C) (Photo Credits: B. Blackman). (D) Average annual precipitation (left, mm) and average annual temperature (right; °C * 10) obtained from the WorldClim BioClim dataset plotted at each population site.

## Methods

### Sample collection, WGS library preparation, resequencing, and variant calling

We obtained genomic short-read resequencing data for 327 individuals from 110 annual populations of *Mimulus guttatus* sampled across Oregon and California [Table S1]. Of these, 237 individuals were newly sequenced for this study; the data for the remaining 90 individuals was initially reported in Colicchio et al., 2021. The methods employed for generating this data and genotype calling are described in the Supplemental Text.

### Population structure

Range-wide population structure was initially examined using Principal Component Analysis (PCA) via PLINK version 1.9 (Purcell et al., 2007). Ancestry assignment was performed using both spatially agnostic and spatially informed methods. We estimated individual ancestries with ADMIXTURE for K = 1 - 10 clusters using default parameters (Alexander & Lange, 2011). The optimal K value was determined by cross-validation [Figure S1]. We also predicted individual ancestries K = 1 - 10 clusters while accounting for physical proximity with TESS3 (Caye et al., 2016), the first step of the ALGATR pipeline (Caye et al., 2016); all inputs were derived from an imputed dosage matrix created using the “vcf_to_dosage()” and “simple_impute()” commands. This matrix is then compared to raster data downloaded from the WorldClim database (Fick & Hijmans, 2017) at 2.5-minute spatial resolution. The optimal K value was selected using the “bestK()” command [Figure S2]. For downstream analyses, we assigned populations to one of the three regional ancestry subsets that emerged from this analysis (Oregon = OR, Northern California = NC, or Central and Southern California = SC) based on the ancestry group with the highest percent affiliation.

To screen for and remove any interspecific hybrids or individuals misidentified as *M. guttatus*, we performed a PCA including our samples and confirmed individuals from *M. laciniatus*, *M. nasutus*, and perennial *M. guttatus* (Ivey et al., 2023). Three putative interspecific hybrid individuals were excluded from downstream analyses.

### Isolation by Distance and Environment

Range-wide signatures of Isolation by Distance (IBD) and Isolation by Environment (IBE) were assessed using tools in the ALGATR pipeline. Multiple matrix regression with randomization (MMRR; Wang, 2013) was performed by running a full model over 99 permutations with seed set = 10 and default parameters. To examine allele frequency turnover patterns along environmental gradients, we performed generalized dissimilarity modeling (GDM) with individual-based sampling and default parameters over 500 permutations (Ferrier et al., 2007; Fitzpatrick & Keller, 2015; Freedman et al., 2010). Methods applied for estimating population genomic parameters are described in the Supplemental Text.

### Climate variables

Climate metrics were obtained from WorldClim (Fick & Hijmans, 2017) and the Basin Characterization Model (BCM; Flint et al., 2021) [Figure S3; Table S2]. We generated BioClim variables (Busby, 1991), a series of 19 monthly climatic combinations from 1970-2000, from WorldClim using the R package sp (Pebesma & Bivand, 2005) at 30 arc-second resolution for our sites.

The BCM balances the hydrologic cycle on a per-raster basis. It integrates temperature and precipitation information with maps of soil depth and porosity and of bedrock geology (Flint et al., 2021; Thorne et al., 2015). The historical climate time series input into the BCM were the PRISM monthly temperature and precipitation values, a gridded dataset covering the continental United States, available at 4 km resolution (Daly et al., 2008). These were downscaled to 270 m resolution using spatial gradient and inverse squared (GIDS) interpolation (Flint & Flint, 2012).

The BCM outputs monthly values for eight variables: snow pack accumulation, snow melt, runoff, recharge, soil moisture (SM), potential evapotranspiration (PET), actual evapotranspiration (AET), and climatic water deficit (CWD). We used the raster outputs from the BCM (Flint et al., 2021; Seymour et al., 2026) to extract monthly time series data from 1970- 2000 for each of our 110 study sites. Water year averages (October through September) were calculated by month for each variable. However, annual averages or totals of PET, AET, and CWD are not effective site descriptors because individual values are calculated dependent on the preceding month’s value. Therefore, we computed the annual Area Under the Curve for these parameters. Because the growing season timing varies greatly across the *M. guttatus* range depending on soil exposure and water availability (Kooyers et al., 2015), we developed coarse metrics corresponding to the predicted growing season start month, end month, and length (Supplementary Text).

To reduce collinearity among subsets of the 39 total BioClim and BCM variables, we collapsed highly correlated variables into groups using the “Cluster Variables” command from JMP Pro 17 software (SAS Institute Inc., Cary, NC). Pairs of variables with an R^2^ value of > 0.7 within their cluster were treated as collinear. To accurately describe cluster variance, we ran PCAs for each cluster of variables separately and extracted the first principal component as the value for the cluster for all sites. Variables with R^2^ < 0.7 across all clusters were retained as single variables. This process yielded 17 climate parameters [Table S3] whose yearly values were converted to z-scores to ensure standardized comparisons across variables.

### Genotype-Environment Association (GEA) analysis

We conducted two types of GEA analysis, one multivariate and one univariate, to identify SNPs with allele frequency patterns associated with variation in environmental parameters. Both GEA analyses were conducted using a pruned vcf where only biallelic SNPs with minor allele frequency > 0.03 were retained. First, we performed partial redundancy analysis (pRDA) (Capblancq & Forester, 2021) using the vegan R package version 2.6-4 (Oksanen et al., 2024). To determine the overall input of the environment alone to genetic variance, we performed multivariate constrained ordination on the first three principal components of variation in our reduced set of 17 environmental variables (envPCs) [Table S4]. To assess how much of the total genetic variation is explained individually by the environment (envPCs), population structure (genPC1 and genPC2), or geographic location (latitude and longitude), we ran a series of three pRDAs, which each include the other two explanatory variables as covariates, and compared these three models to a final model that examines all three factors together. For all further pRDA models, we use the environmental pRDA, which includes environment as an explanatory variable and includes population structure and geographic location as covariates.

The pRDA models were run on the whole dataset (full range, or FR) and also separately run for subsets of individuals corresponding to each of the three regional ancestry groups (OR, NC, and SC) as determined by TESS3. Population structure data for each regional ancestry group was subset from the full range genPC1 and genPC2 values to ensure consistent comparison of axes. Outlier RDA value cutoffs were set to the Bonferroni correction threshold for each model (FR: -log_10_(p) = 7.409; OR: -log_10_(p) = 7.205; NC: -log_10_(p) = 7.319; SC: -log_10_(p) = 7.324).

Second, we ran Latent Factor Mixed Models (LFMMs) with the vegan R package (Oksanen et al., 2024) using the *ridge* method for its increased ability to handle multicollinear data (Caye et al., 2019). The number of latent factors was determined using the *elbow* method for each model (best K estimation by model: FR = 6; OR = 3; NC = 6; SC = 4) [Figure S2]. Using the lfmm_ridge() command, we ran all four univariate constrained ordination models using the respective latent factors as covariates for each of our 17 clustered climate variables. Outlier LFMM Wald test value cutoffs were set at the Bonferroni correction threshold for each model (FR: -log_10_(p) = 7.408551; OR: 7.205; NC: 7.319; SC: 7.324).

We used Weighted-Z analysis (WZA) approach to enhance signal from noise by coalescing information from multiple nearby SNPs within fixed windows of 200 SNPs to better detect environment-associated haplotypes (Booker, 2024). This window-based outlier identification approach was run on the p-value output from each of our RDA and LFMM models, and outlier congruence across methods was reviewed as a corroborating indicator of true positives. Windows with WZA p-values < 0.05 were considered outliers. Our final lists of GEA outliers consist of SNPs which both pass their respective GEA method and WZA significance thresholds listed above and were not located in intervals flagged as likely multicopy regions by ParaMask (Tjeng et al., 2025).

### Genome-Wide Association (GWA) analysis

To explore whether any GEA hits explain heritable trait variation, grew the subset of 237 individuals newly resequenced in this study in a common garden and measured ten relevant phenotypes. Methods for phenotype measurement and GWA analysis are described in the Supplemental Text.

### Focal outlier Selection

Each identified outlier SNP was further evaluated for gene identity, mutational impact, and GO term enrichment as a measure of relative importance. First, for each outlier SNP, we extracted the gene code ID(s) and description(s) for the *M. guttatus* var. IM62 v3 reference genome annotations which fall within gene annotations with Phytozome’s BioMart tool (https://phytozome-next.jgi.doe.gov/biomart/). Then, the best hit *Arabidopsis thaliana* TAIR 10 annotation (https://phytozome-next.jgi.doe.gov/info/Athaliana_TAIR10) was identified by blastp (Altschul et al., 1990). To prioritize GEA or GWA outliers for downstream analysis, each SNP was ranked using an ordinal stratification method to combine significance rankings across methods. The SNPs that scored collectively highest across both metrics (RDA/LFMM, or LMM and BSLMM) are thus ranked highest. Finally, the predicted impacts of SNPs were assessed using snpEff and snpSift (Cingolani, Patel, et al., 2012; Cingolani, Platts, et al., 2012).

Phytozome’s PhytoMine tool (https://phytozome-next.jgi.doe.gov/phytomine/) was used to to retrieve annotations and GeneOntology (Ashburner et al., 2000); https://geneontology.org/) and then calculate GO term enrichment for each list of outlier SNPs using a False Discovery Rate correction and a minimum adjusted p-value of 0.05.

## Results

### Regionally subdividing GEA analysis increases the fraction of genetic variation specifically explained by environmental variation

We first assessed patterns of population structure across the *M. guttatus* range and its relationships to geography and climate. Both ADMIXTURE (non-spatial) and TESS3 (spatial) ancestry estimation methods yielded optimal values of K = 3 groups, roughly corresponding with three latitudinally disjunct regions: Oregon (OR), Northern California (NC), and Central and Southern California (SC) [Figure 2a,b]. Consistent with these groupings, the first principal component of genetic variation (genPC1) is associated with latitude (R^2^= 0.148; p<0.0001) [Figure 2c]. Individuals were assigned to an ancestry group based on the TESS output (FR: n = 327; OR n = 117; NC: n = 80; SC: n = 130). Nucleotide diversity (π) values across the full range (FR) and within ancestry groups are similar (FR: 0.0194; OR: 0.0163; NC: 0.0193; SC: 0.020), and low population pairwise F_ST_ values suggest that most genetic diversity is shared among regions (OR/NC: 0.044 ± 0.080 [s.d.]; NC/SC: 0.033 ± 0.104; OR/SC: 0.056 ± 0.067).

**Figure 2:**
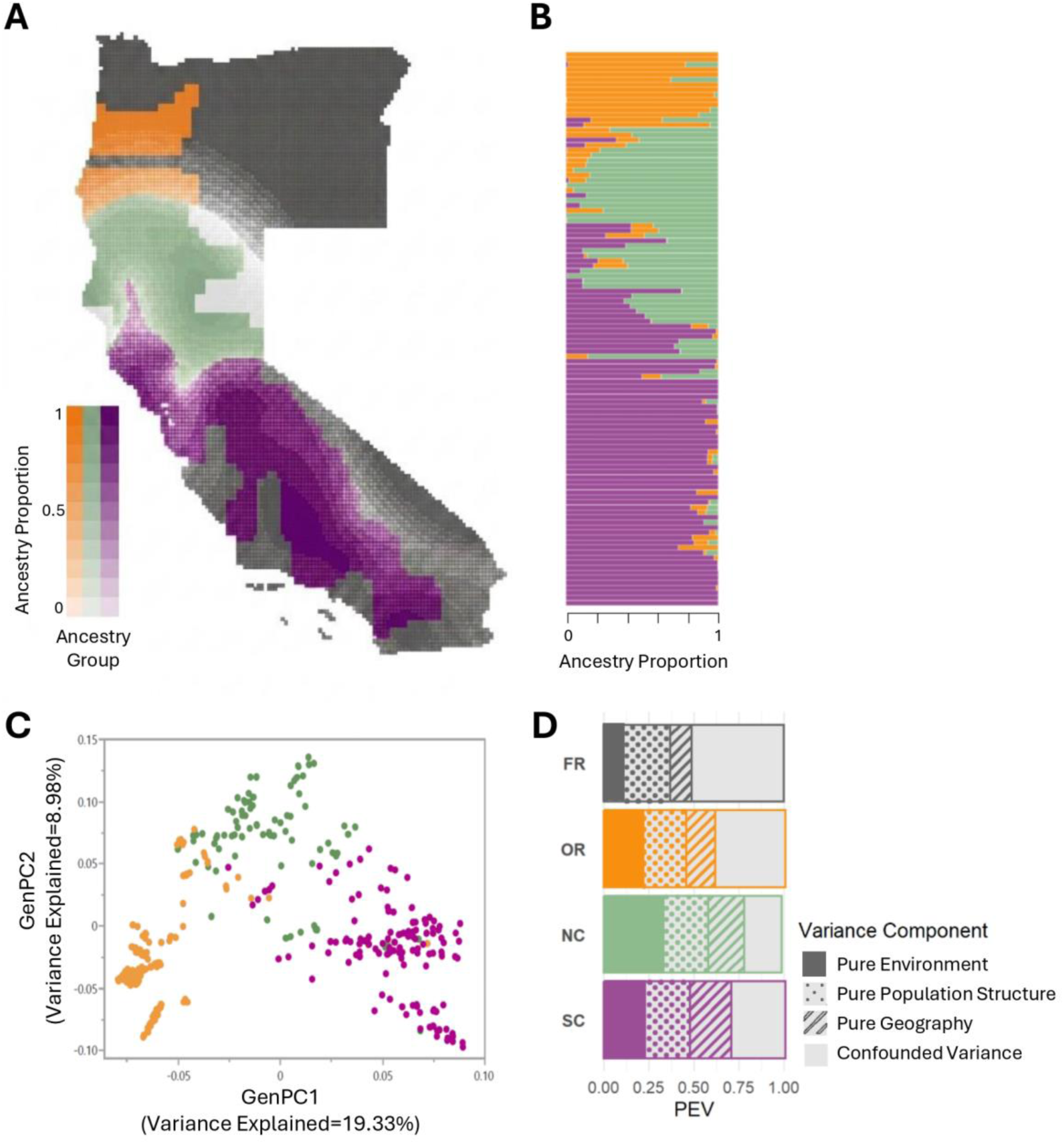
*M. guttatus* Resolves into Three Ancestry Groups Largely Correlated with Latitude. (A) TESS3 population site ancestry majority group identity projected onto raster-level data and colored by TESS3 ancestry grouping. (B) ADMIXTURE population site ancestry arranged by latitude ascending from left to right and colored by ADMIXTURE ancestry grouping. (C) GenPC1 and GenPC2 scores colored by TESS3 ancestry grouping. All plots are colored by ancestry group assignment: Oregon (OR) = orange, Northern California (NC) = green, and Central and Southern California (SC) = purple. (D) Variance partitioning of the amount of explainable genetic variance attributable to each included metric as output by the full-range (FR) model and the three regional pRDA models.

Previous work has found that genetic differentiation increases with geographic and environmental distance in *M. guttatus* and related taxa (Sexton et al., 2016; Twyford et al., 2020), and we observe a strong association between genPC1 and the first principal component of total environmental variation (envPC1) in our dataset (R^2^ = 0.652, p <0.0001) [Table S5; Table S6]. Therefore, we conducted formal tests for isolation by distance (IBD) and isolation by environment (IBE). An MMRR analysis revealed patterns of both IBD (β_geo = 0.57, *p* = 0.01) and IBE (β_envPC1 = 0.53, *p* < 0.001) [Table S7; Fig. S4]. To account for potential nonlinear relationships between genetic and environmental distances, we also performed generalized dissimilarity modeling (GDM). Total environmental dissimilarity across all three envPCs combined explains 44.3% of the total deviance in genetic composition (R^2^ = 0.443; p = <0.001) [Figure S4; S5; S6]. Thus, climate variation has shaped patterns of genetic variation across sites in *M. guttatus*.

In light of these findings and because environmental distance and geographic distance are also correlated with each other in our dataset [Table S7], we performed pRDA to jointly quantify the contributions of population structure, geographic distance, and environmental factors (envPC1–3) to genetic variation. Of the 18% of the total genetic variance explainable by the pRDA model, 10% is uniquely attributable to environmental terms, and 27% is uniquely attributable to population structure [Figure 2d, Table S8]. Roughly half of explainable genetic variance could not be associated with any individual predictor but rather is confounded among predictors, consistent with the correlations among these predictors described above. As compared to results from pRDA models that include either just BioClim or just BCM environmental parameters, a lower proportion of the explainable genetic variance is confounded among predictors in the full model that includes environmental parameters derived from both BioClim and the BCM [Table S9]. Thus, these results highlight the value of incorporating both climatic and hydrological variables when evaluating how environmental factors relate to patterns of genetic variation.

Because genPC1 is associated with affiliation to the three ancestry groups [Figure 2c], we hypothesized that the confounded variance component would be lower and the environmentally explainable variance would be higher for pRDAs conducted separately by regional subset. Indeed, the proportion of genetic variation explained solely by environmental factors increased by two- to three-fold in all three regional models (OR: 22%; NC: 33%; SC: 23%), largely reflecting reductions in confounded variance [Figure 2d; Table S8]. These results demonstrate that subdividing range-wide datasets into regional groups prior to GEA analyses can improve inference in taxa where major components of population structure align with environmental gradients. Doing so appears to strike a balance between removing false positives caused by population structure and false negatives due to its collinearity with the environment and region-specific adaptive variants.

### Environmental Heterogeneity Results in Distinct Selective Pressures across Regions

Regional targets of local adaptation may be expected to differ when the agents of selection or their range breadth differ by region (Le Corre & Kremer, 2012). Therefore, we first investigated the degree to which our three ancestry groups inhabit distinct climates. EnvPC1 is significantly different among all regional pairs [Figure S7a], and 95% confidence interval ellipsoids showed minimal environmental overlap between OR and SC sites [Figure 3a], indicating that the three ancestry groups experience distinct climate regimes. Consistent with this finding, across three separate regional PCA models, the environmental variables contributing most to envPC1, envPC2, and envPC3 of each regional model are largely unique [Figure S8]. For example, while precipitation (Group6) contributed most to envPC1 across the full range, Snow (Group4), Runoff, and Evaporation were the largest contributors to envPC1 for OR, NC, and SC, respectively. Across the top three envPCs from the four models, 10 of the 12 most contributory environmental variables are unique, with only Elevation and Runoff appearing twice. These patterns indicate that populations from different ancestry groups experience distinct environmental regimes with divergent axes of variation and likely different primary selective pressures.

**Figure 3:**
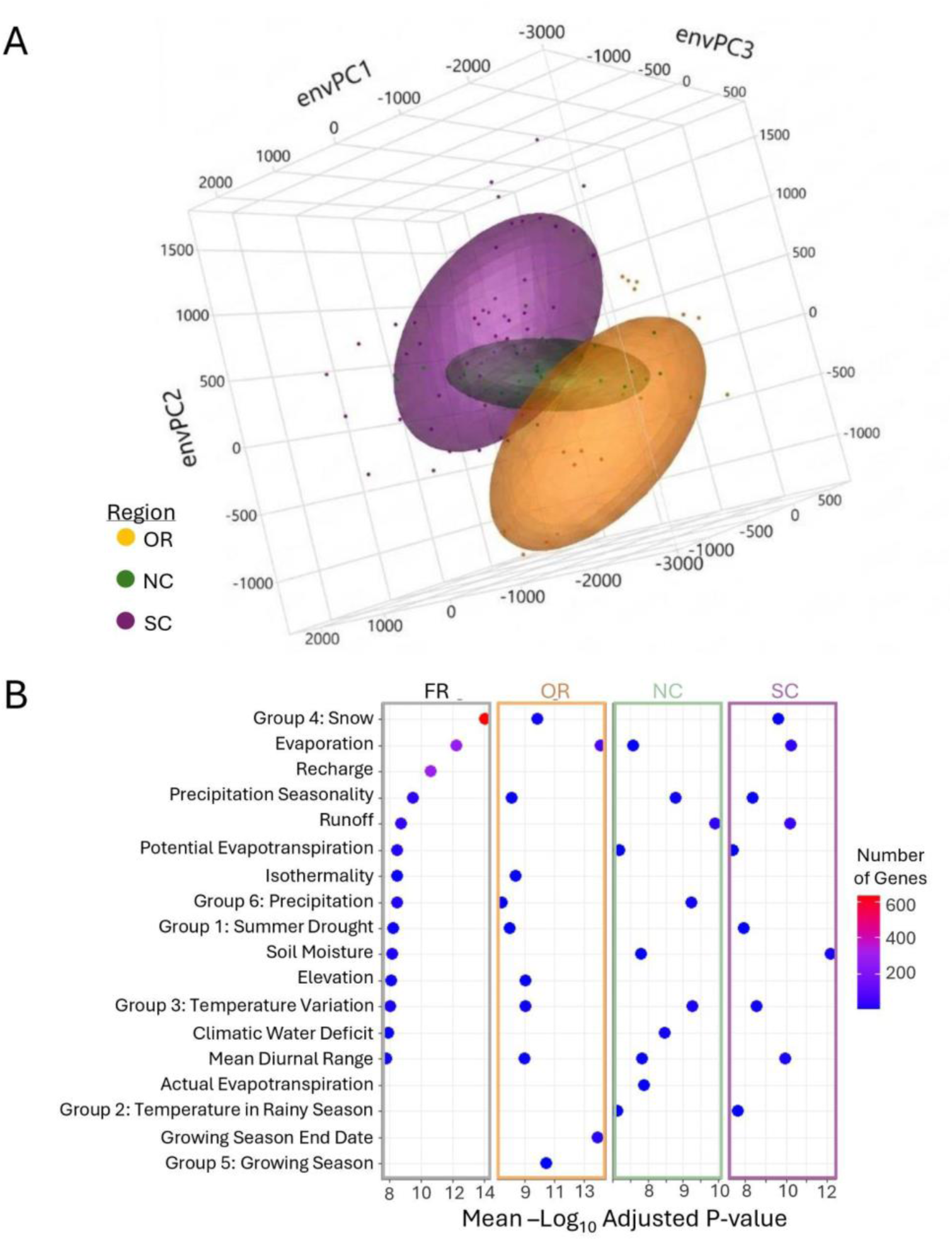
Environmental Variables of Importance Vary Across Regional Models. Models are denoted as: full range (FR), Oregon (OR), Northern California (NC), and Southern California (SC). (A) Multi-dimensional depiction of environmental variance across regions using the first three environmental principal components. (B) The mean -log10 adjusted LFMM *P*-value for all SNP outliers for each environmental variable. Values are colored by the number of genes included in each variable outlier set. Variables are ordered by the mean -log 10 adjusted *P*-value in the full range model.

To assess whether these three distinct environmental landscapes impose different selective pressures, we first identified the environmental variables contributing most to the largest axis of variation in our RDA models [Figure S8, S9; Table S10]. For the FR model, Elevation contributed most to envPC2, which in turn loaded most strongly on RDA1, explaining 41% of total variance. Regionally, Group4 (Snow) in OR, Group2 (Temperature during the Rainy Season) in NC, and Elevation in SC contributed most to their respective RDA1 values, indicating that variation in each region is shaped by distinct environmental pressures.

We also evaluated whether regions differ in which agents of selection are most important by comparing which environmental variables using the output of LFMM analyses. Specifically, we evaluated which individual parameters stood out for two possible proxies for importance as an agent for selection: number of outlier SNPs and the mean p-value for the outlier SNP sets.

For the FR data, we find that Group4 (Snow), Evaporation, and Recharge stand out as agents of selection [Figure 3b]. Again, the relative importance of environmental variables differs across each regional subsect. At the regional scale, Growing Season Timing appears most important in OR, Precipitation in NC, and Soil Moisture in SC [Figure 3b]. Together, this evidence indicates that climate factors vary substantially in their importance as agents of selection across the range. Therefore, we predicted that local adaptation to climate would involve more unique than shared genetic targets of selection at a regional scale. Nonetheless, some shared targets of selection may be expected for any environmental conditions that have similar regional gradients or where similar phenotypic responses have been favored to evolve in response to different selective pressures.

### Targets of Selection are Largely Distinct Across Regions

We next applied two GEA approaches, pRDA and LFMMs, to examine how extensively the genomic regions putatively contributing to climate adaptation overlapped among the range-wide model and each of three region-specific models. We first summarize the results of this array of individual multi- and univariate models before then examining the amount of congruence across approaches and regions to test our predictions. The FR pRDA model detected 760 outlier SNPs corresponding to 251 genes [Table S11]. At the regional scale, the OR pRDA detected 473 outlier SNPs (139 genes), the NC pRDA detected 300 SNPs (44 genes), and the SC pRDA identified 164 SNPs (81 genes) [Table S11]. To better leverage shared signals of adaptive differentiation among sites closely linked to causal variants, we processed the pRDA results using Weighted Z-Analysis (WZA; Booker, 2024), highlighting genomic windows that are significant GEA hits. This analysis detected 214 significant windows for the FR pRDA model, 112 significant windows for the OR pRDA model, 41 significant windows for the NC pRDA model, and 72 significant windows for the SC pRDA model [Figure S10, S11; Table S11].

The FR LFMMs detected 5,183 SNP-environment associations (1,112 genes) across the 17 environmental variables tested and 751 significant windows [Table S13]. Each set of 17 regional LFMMs detected fewer SNP-environment associations (OR: 194 SNPs across 102 genes; NC: 118 SNPs across 57 genes; SC: 179 SNPs across 97 genes) and significant WZA windows (OR: 98 windows; NC: 68 windows; SC: 105 windows) than the FR model [Table S13].

Many significant GEA hits were detected with both the RDA and LFMM approaches. Across all models, 134 SNPs in 70 genes were identified by at least one RDA and one LFMM model [Table S14]. Within the FR model, 131 SNPs spanning 68 genes overlap between methods, with the majority associated with Group4 (“Snow”) (120 SNPs in 60 genes) and Recharge (30 SNPs in 16 genes). [Table S14].

We find limited overlap among environmentally associated genomic variation across regional models [Figure S12]. Of 6,405 genomic windows (200 SNPs each), 1,008 and 1,022 outliers were detected in RDA and LFMM, respectively [Table S11, S13]. Only 48 pRDA windows and 44 LFMM windows overlap across regions [Figure 4a,b], with the majority of windows unique to their respective region. Likewise, we find little overlap across regions in the individual genes associated with environmental variation. We identify 50 genes (RDA; 10.8%) and 31 genes (LFMM; 2.3%) with outlier SNPs in common across at least two regions [Figure 4b, S11, S13]; outlier genes most frequently overlap between the FR and SC models. The majority of these overlapping genes are associated with Group4 (“Snow”), the key variable for the FR dataset [Figure 3b, Table S13]. On a single-variant scale, overlap is similarly low: 80/1,364 SNPs (5.8%) in RDA and 15/5,068 SNPs (0.2%) in LFMM overlapped across regions, with no SNPs shared among more than two regions [Figure S14; Table S13]. Regardless of the genomic scale, the vast majority of hits are region-specific, indicating that adaptive differentiation in *M. guttatus* often occurs along local-scale environmental gradients and that the genetic basis of climate adaptation is not fully reflected by range-wide modeling.

**Figure 4:**
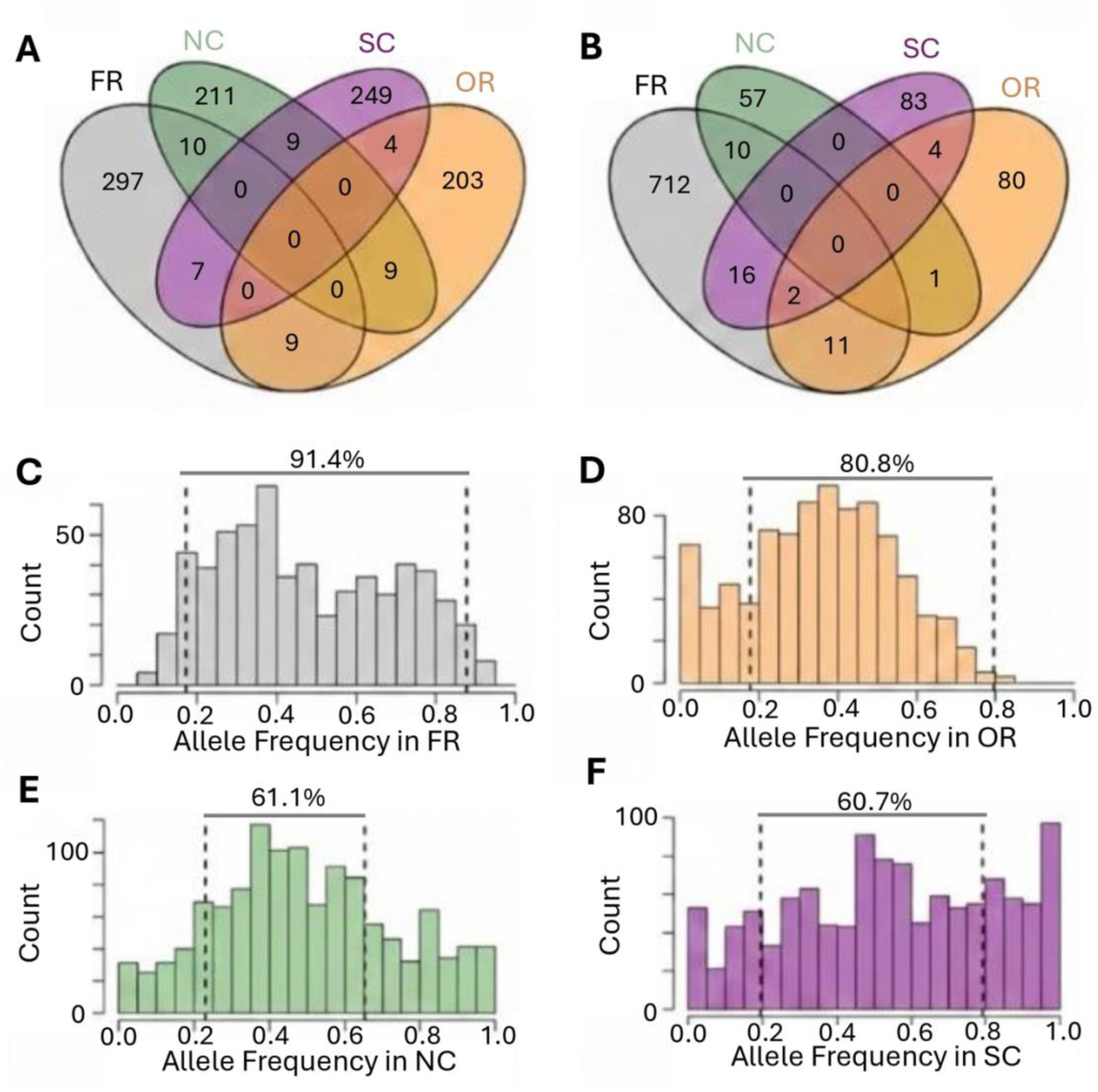
RDA Hit Disparities Across Regions Are Not Due to Lack of Available Genetic Variation. Venn diagram showing the overlap of environment-associated WZA windows (p value < 0.05) across (A) RDA regional models or (B) LFMM regional models. C-F) Allele frequency spectra of the RDA hits which appear in any of the other regions but do not appear in as hits in the respective focal region. Detectability thresholds (dashed vertical lines) for genotype-environment associations are defined by region as the minimum and maximum allele frequencies observed for significant hits in the focal region. The percentage of the total variants that fall within the detectability thresholds is displayed between the thresholds.

### Regional differences in standing variation largely do not explain GEA hit incongruence

Although distinct selective pressures may underlie the limited congruence in targets of selection among regions, methodological factors, including reduced statistical power at smaller scales, differing overlap with population structure, or variation in segregating polymorphism may also contribute (Coop et al., 2010; De Mita et al., 2013; Garant et al., 2007; Lotterhos, 2023). To assess whether the absence of signal in a specific region reflects limited genetic variation, we examined RDA hits present in at least one other region but absent in the focal region. If one allele at most of these RDA hits is only present at low frequency in the region lacking the signal, incongruence might stem from insufficient variation. Notably, for the majority of our RDA SNP hits that were absent in a focal region but found in at least one other, both alleles segregate at intermediate frequencies [Figure 4c-f], indicating that ample genetic variation exists at these sites in each region. When quantified accounting for differences in sample size using the highest and lowest allele frequencies among the GEA hits in a focal region as proxies for the bounds for statistical detectability in that region, we observe that >60% of SNPs detected as outliers in at least one other model but not in the focal region still fall within the expected detectability range across all models. [Figure 4c-f]. These results suggest that regionally private standing variation for potentially adaptive variants is not the predominant source of regional incongruence.

We investigated whether the absence of shared standing genetic variation or differences in agents of selection may explain regional incongruence using a case study of a prominent gene. We examined a homolog of *TBP-associating Factor 5* (*TAF5*), a subunit of the TFIID and SAGA complexes essential for plant development and abiotic stress responses (Mougiou et al., 2012). *TAF5 is* among the top hits in the FR and OR RDA models but lacks associations in the NC and SC RDA models [Table S11]. Among the 43 intronic or synonymous SNPs significant in the FR model, the OR model, or both, the most significant SNP (Chr_14:3126435; Full Range RDA p = 6.56E-18; WZA p = 4.03E-06) [Table S11] is an iHS outlier (iHS = 1.337), and shows weak but extended haplotype homozygosity (EHH) and linkage disequilibrium decay over ∼5 kb [Figure S15], well above the genomic average of ∼1 kb [Figure S16]. *TAF5* also contains more SNPs detected by both the RDA and LFMM than any other gene, with all 125 overlapping SNPs in the Full Range model associated with Group4 (“Snow”) or Recharge.

If differences in standing genetic variation explain the incongruent detection of *TAF5* GEA hits, we would expect few of these SNPs to be at intermediate frequencies in NC and SC. Yet, while some *TAF5* SNPs do have low minor allele frequencies in NC and SC, many still segregate at intermediate frequencies in these regions [Figure 5a]. Notably, nearly all GEA hits cluster around 0.5 in OR, including the most significant FR model SNP, Chr_14:3126435, and most OR individuals are homozygous at this SNP, with a deficit of heterozygotes [Figure 5d].

**Figure 5:**
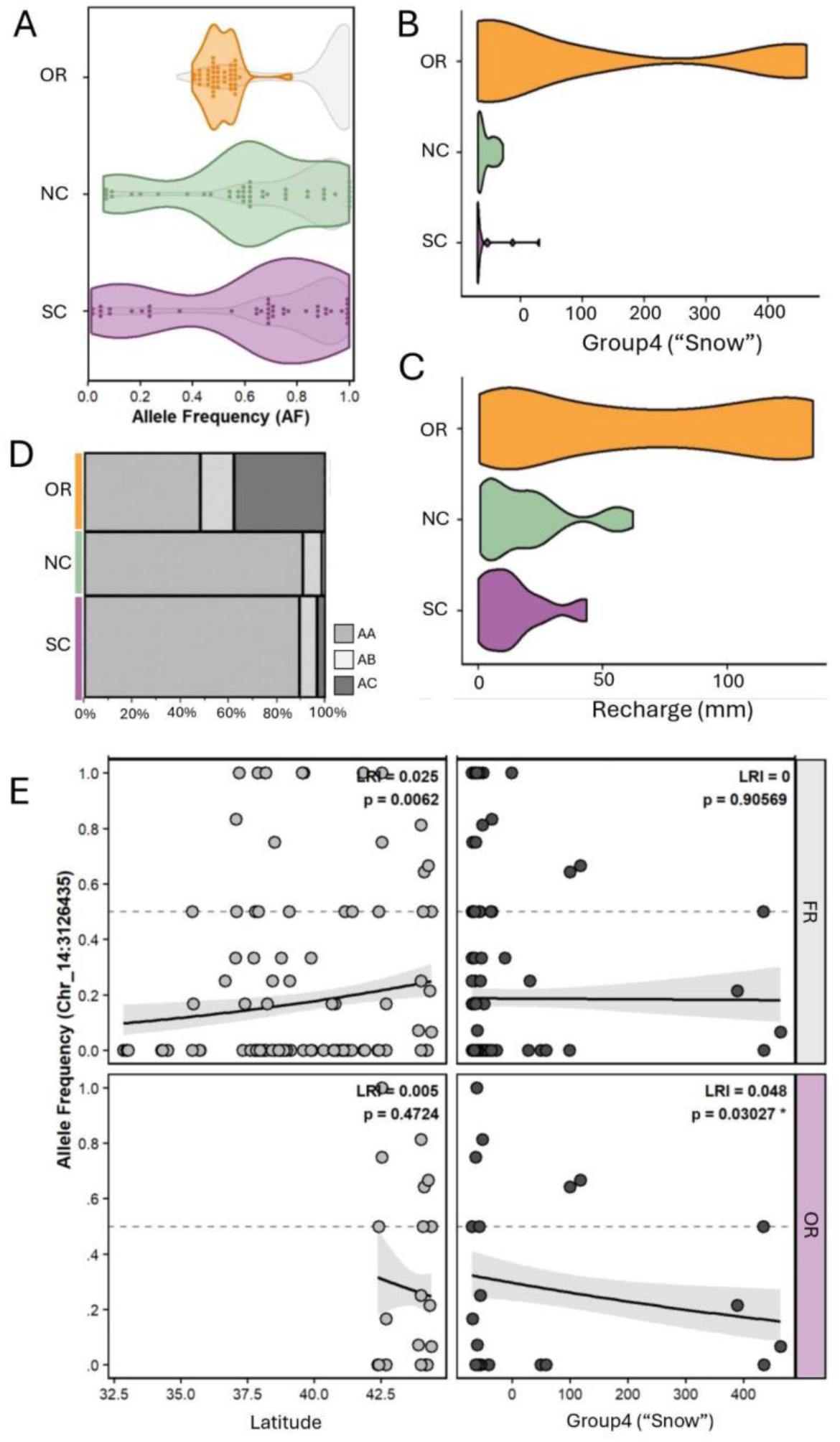
Demographic and Environmental Factors Result in Incongruence across Regions. (A) Allele frequencies of all SNPs within *TAF5*. Significant GEA SNPs are visualized as colored points and their associated colored violin plots (orange = OR, green = NC, and purple = SC). Total SNPs which fall in *TAF5* are visualized as the underlying grey violin pilots for each region. (B) Violin plots visualizing environmental range for Group4 (“Snow”) in mm across regions. (C) Violin plots visualizing environmental range for Recharge in mm across regions. (D) Genotype frequencies by region at the most significant TAF5 GEA hit [Chr_14:3126435] (A = reference allele, B = alt allele). (E) Clinal associations with Latitude and Group 4 (“Snow”) modeled at the individual level across two spatial windows (FR and OR) using weighted binomial generalized linear models (glm). Predictive power was quantified via McFadden’s Likelihood Ratio Index (LRI) relative to an intercept-only null baseline model, with final logistic sigmoid trajectories visualized over aggregated population allele frequencies. P-values were derived directly from the regression slope coefficient summaries.

Although this SNP segregates within NC and SC, most individuals are homozygous for the major allele. These patterns are consistent with maintenance of the polymorphism by region-specific divergent, spatially varying, or temporally varying selection and not by overdominance.

If differences in climatic selection explain the incongruent detection of *TAF5* GEA hits, we would expect to find that the environmental factor most associated with *TAF5* SNPs in OR has a disparate pattern of variation in NC and SC. Consistent with this possibility, we find that Oregon has substantially more breadth for the two environmental variables associated with *TAF5* SNPs in our LFMM models, Group4 (“Snow”) and Recharge [Figure 5b,c]. We also note that the environmental factors most predictive of allele frequency at Chr_14:3126435 depend on scale [Figure 5e]. Across the full range, its allele frequency is associated with latitude, but not specific climate factors like Snow (Group4) [Figure 5e]. Conversely, within Oregon, its allele frequency is significantly associated with Snow but not Latitude [Figure 5e]. This scale-dependent reversal demonstrates how broad geographic clines may obscure the signal of local associations between genotype and particular environmental factors.

### Overlapping GEA signals highlight flowering time regulation, light signaling, and ABA signaling as common targets of selection

While GO analysis showed enrichment for relatively general functional categories in results of the FR RDA and LFMM models [Table S12], the subset of GEA hits that overlapped across multiple models highlighted several specific signaling pathways as frequent contributors to climate adaptation. For instance, several hits occurred in genes homologous to regulators of the *FLOWERING LOCUS C* and *MADS AFFECTING FLOWERING* (*FLC/MAF*), which are involved in temperature-dependent regulation of flowering time [Table S11, S13] (Airoldi et al., 2015; Scortecci et al., 2001; Suter et al., 2014; Werner et al., 2005). Multiple GEA hits were detected for homologs to components of the FRIGIDA (*FRI*) complex or the chromatin modifying complex the FRI complex recruits to activate *FLC* (Choi et al., 2011). These signals include multiple hits in two homologs of *FLC EXPRESSOR* (*FLX*; Lee & Amasino, 2013) associated with Snow in the FR LFMM and Growing Season End Date in the OR LFMM, respectively; and *FRIGIDA-ESSENTIAL 1* (*FES1*; Schmitz et al., 2005) associated with Elevation the FR RDA model and with Group4 (“Snow”) in the FR LFMM model. The most significant GEA hit in a homolog of *SUPPRESSORS OF MEC-8 AND UNC-52 1* (*SMU1*) [Chr_03:23498841], a spliceosome protein involved in *FLC* repression (Long et al., 2024), is associated with Snow in the FR LFMM and Recharge in SC LFMM [Figure S17; Table S13], exhibits extended LD (∼100 kb) [Figure S16], and is an outlier for EHH (∼20 kb) [Figure S17]. Although *SMU1* also functions in response to magnesium stress (Feng et al., 2020), the convergence of GEA analyses on epigenetic regulators of *FLC/MAF* genes strongly indicates that these mechanisms harbor ample variation that has been targeted by natural selection in *M. guttatus*.

We also detected patterns implicating light signaling pathways in climate adaptation. Phytochrome E (phyE) contains the most significant hits in the FR and SC pRDA models and has variants associated with Evaporation across FR, OR, and SC LFMM models [Table S11, S13]. The most significant full-range pRDA hit (Chr_08:18325023) in PhyE is an exonic polymorphism that exhibits high LD and EHH (∼2 kb) [Figure S18]. Variation in phytochrome A (phyA) is also associated with Evaporation in the FR and SC LFMM models and bears the third most significant SNP across all SC LFMMs combined [Table S13]. This SNP, Chr_02:5130164, is an upstream variant that shows extended LD (∼100 kb) and is an EHH outlier (∼3 kb) [Figure S19]. A homolog of *SUPPRESSOR OF PHYA-105 1* (*SPA1*), which targets phyA for degradation (Hoecker et al., 1999; Zhu et al., 2008), is FR RDA outlier and a FR LFMM outlier associated with Group4 (“Snow”) [Table S14]. Together, these results suggest that differentiation that light-regulated developmental processes like germination and flowering may be key targets of selection.

We also found concordant hits across pRDA and LFMM approaches for many ABA-signaling components that could relate to root development and stomatal function [Table S11; Table S13]. A homolog of *YODA, a* MAPKK kinase involved in auxin-dependent post-embryonic root development and ABA signaling (Lukowitz et al., 2004; Smékalová et al., 2014; H. Wang et al., 2007), is the most significant NC pRDA hit and the 19th most significant FR pRDA hit, with 7 LFMM SNPs associated with Snow (Group 4), Recharge, or both. Other ABA-related hits include *ABA-OVERLY SENSITIVE 1* (*ABO1*), associated with Temperature in the Rainy Season and the most significant FR LFMM locus (An et al., 2016), and *MIZU-KUSSEL 2*, the fourth most significant hit for Soil Moisture in SC (Guo et al., 2014; Miyazawa et al., 2009; Pang et al., 2023). The stomatal-regulation gene *OMISHA* (Linster et al., 2015) is the only locus overlapping across three pRDA models and associates with Elevation and Snow (Group 4) in the RDA and Recharge in the FR LFMM.

### GWA Analysis Also Highlights Positive and Negative Regulators of FLC/MAF genes

To assess whether any GEA hits may explain locally adaptive trait variation, we conducted GWA analyses for 10 phenotypes. Because trait data were available for only a subset of resequenced individuals (n = 237), including few Oregonian samples, we did not subdivide these analyses by region. Consistent with previous work (Kooyers et al., 2015), most of the phenotypes are distributed in clines along environmental gradients [Table S15]. All phenotypes measured except Peduncle Length displayed high heritability (PVE > 0.5), and several traits are influenced by one or more large effect loci (PGE > 0.4) [Figure 6a]. Thus, most the traits measured have histories of adaptive differentiation relevant to the climate factors used in our GEAs, with much of this variation being heritable.

**Figure 6:**
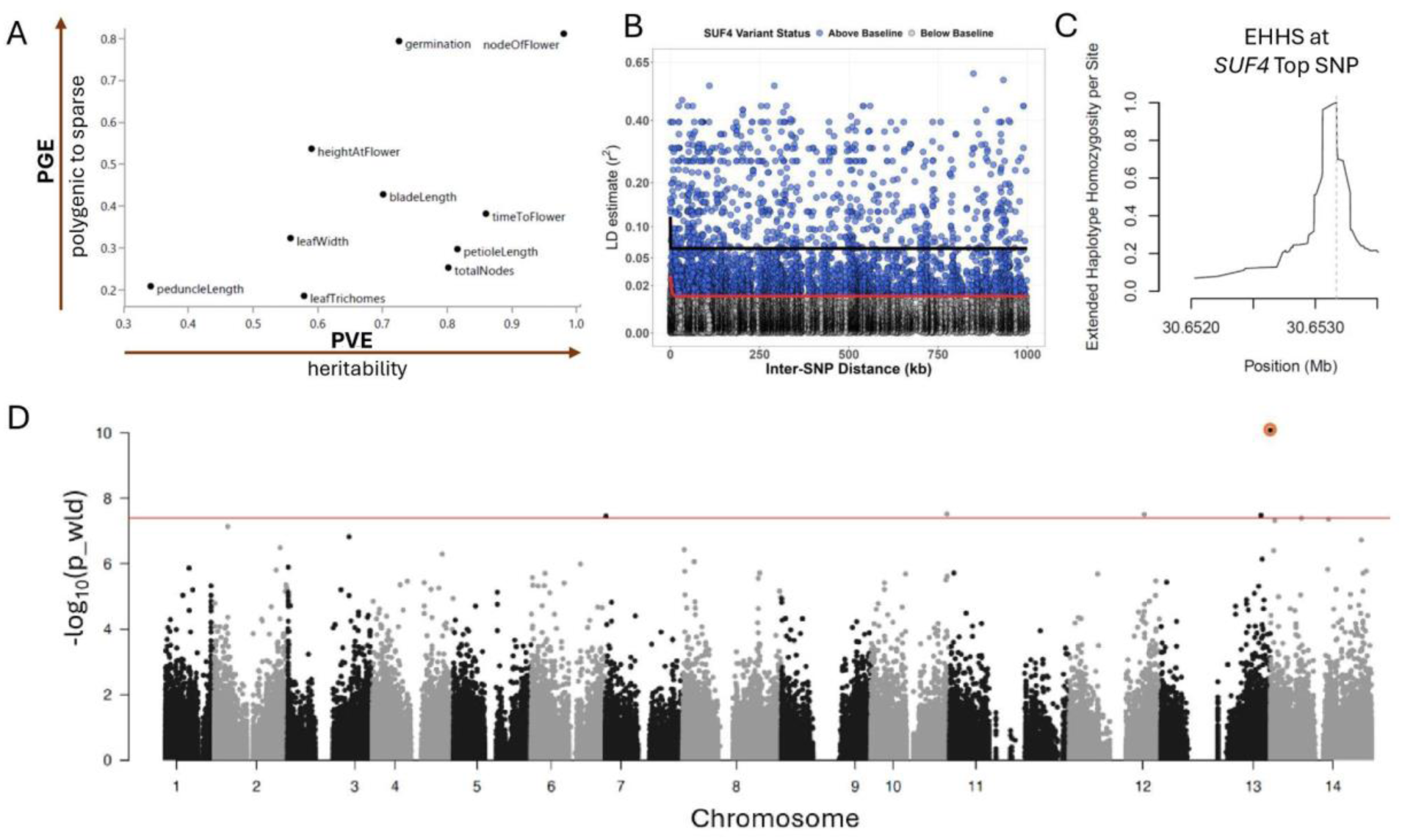
GEA hit in *SUF4* is Associated with Height at Flowering and Signatures of Selection. (A) Proportion of Variance Explained (PVE), a measure of heritability, and Proportion of Genetic Variance Explained (PGE), a measure of polygenicity, as estimated by BSLMM for ten traits. (B) FR linkage disequilibrium (LD) decay plotted against the distance between *SUF4* locus variants and nearby SNPs. Red line = Background genome-wide decay. The solid black curve tracks the extended LD threshold specific to the elevated *SUF4* candidate sweep variants highlighted in blue. (C) Full Range per site Extended Haplotype Homozygosity (EHH) around the most significant non-synonymous SNP in *SUF4*. (D) Whole genome Manhattan plot of uncorrected –log10 Time to Flower LMM p_wld values. The most significant non-synonymous SNP [Chr_13:30653171] is circled in orange. Red line = Bonferroni correction significance threshold.

Across the 10 phenotypes, GWA mapping detected 116 significant SNPs corresponding to 93 genes [Table S16]. Only one SNP, Chr_04:21534507, is a significant hit in both RDA and GWA analyses [Table S11, S16]. It is associated with Petiole Length, is downstream to the chlorophyll biosynthesis regulator *FAR1-RELATED SEQUENCE 5* (K. Kim & Choi, 2025), and is in linkage disequilibrium with SNPs up to ∼100kb away [Figure S20; Figure S21a]. We found more overlap between the GWA and LFMM hits, with 14 of the 16 overlapping SNPs coming from the FR LFMM hit set [Table S17]. Notably, many of these SNPs are in flowering time regulators. The most significant GWA SNP across all 10 phenotypes [Chr_13:30653171] falls upstream to a homolog of *SUPPRESSOR OF FRIGIDA 4* (*SUF4*), is associated with both Evaporation and Time to Flower, and is in a genomic region with extended EHH (∼2kb) over which linkage disequilibrium decays slowly (∼100kb) [Figure 6cd, S21; Table S17]. In *Arabidopsis*, *SUF4* forms a complex with *FRI* and acts as a transcriptional activator of *FLC* in a temperature-dependent manner (Choi et al., 2011; S. Y. Kim & Michaels, 2006; Meyer et al., 2025; Yang et al., 2023). Notably, significant GWA hits are also found in another member of the *FRI* complex, *VERNALIZATION INDEPENDENT 5* (Height at Flowering, 8^th^-most significant hit across all traits) (Greb et al., 2007; He et al., 2004), and in additional regulators of *FLC* mRNA abundance, including *EARLY FLOWERING IN SHORT DAYS 3* (Height at Flowering, second-most significant LMM hit overall) and *ABSCISIC ACID HYPERSENSITIVE 1* (Node of First Flower, most significant BLSMM hit overall) (Bezerra et al., 2004; S. Y. Kim et al., 2005).

Together, the GWA results particularly highlight regulators of *FLC/MAF* gene family members as harboring genetic variation for phenological traits in *M. guttatus*.

## Discussion

Our results highlight the complexity associated with determining the agents and targets of selection underlying climate adaptation. We find evidence of genome-wide population structure associated with both geographical and environmental gradients, and climate adaptation across the full range is highly polygenic. However, parsing the range-wide dataset into separate regions provided considerable gains from the full-range models. Regional GEA models increased the fraction of explainable genetic variation specifically attributable to environmental variation, revealed how the most prominent agents of selection substantially differed across range-wide and regional models, and detected environment-associated variants that are largely non-overlapping between regions. The differences among regional models are not primarily due to differences in standing genetic variation or regionally private alleles, but are likely associated with differences in climate breadth between regions. Despite these differences, genomic variation associated with reproductive timing and response to water stress appear important across regions. Below, we discuss our results in the context of the evolutionary genomics of local adaptation and in predicting responses to changing climates.

### Accounting for regional population structure improves capacity to quantify environmental contribution to patterning of genetic variation

We find that genetic variation in annual *M. guttatus* populations is patterned by IBD and IBE, reflecting a complex history of migration and climate adaptation. The annual populations sampled group into three regional genetic ancestry clusters structured by latitude (Figure 2), mostly consistent with previous results that have often included a broader sampling of the *M. guttatus* species complex (Frayer et al., 2026; Ivey et al., 2023; Kooyers et al., 2015; Twyford & Friedman, 2015). For instance, the OR and SC clusters largely correspond to the northern and southern *M. guttatus* clades defined in previous phylogeographic analyses. However, the NC cluster represents a somewhat novel grouping that best corresponds to a clade recently defined as including *M. glaucescens* and northern Sierran *M. guttatus* populations (Frayer et al., 2026; Ivey et al., 2023). In our sampling, which includes many newly resequenced collections from this area, this cluster covers an area extending well beyond the northern Sierra Nevada into the Klamath and Trinity ranges. We will be interested to learn whether this grouping persists and helps resolve alternate biogeographic hypotheses about the post-glacial history of this species when our data are integrated with representatives from throughout the *M. guttatus* species complex (Twyford et al., 2020), and how these findings then relate to broader biogeographic patterns observed across plant taxa also included in the California Conservation Genomics Project (Shaffer et al., 2022). Likewise, we will be interested to learn whether reproductive barriers observed between Sierran and Northern *M. guttatus* populations are also found for non-Sierran NC populations or are specific to the Sierra (Frayer et al., 2026).

We find that patterns of population structure, geography, and environmental variation are all correlated with one another, creating a significant challenge to attributing allele frequency patterns to specific environmental factors. Patterns of IBD and IBE are consistent with previous results (Twyford et al., 2020), and temperature largely decreases and precipitation largely increases with latitude along the West Coast (Dettinger et al., 1998). Differentiation in the timing of seasonal flowering is important for local adaptation and may be one key factor that promotes IBE in *M. guttatus* (Friedman & Willis, 2013; Hall & Willis, 2006; Kooyers et al., 2015, 2019).

Given this challenge, we assessed whether subdividing the full range sample into regional groups defined by ancestry inference prior to pRDA modeling would allow a greater proportion of the explainable genetic variance to be attributed to individual factors rather than to variance confounded by multiple factors. This approach was successful. In all three regional models, the proportion of explainable genetic variance attributable to environmental factors alone increased compared to the full range model, consistent with prior studies that found improved resolution of genotype-environment associations when controlling for hierarchical structure (e.g., (Forester et al., 2018; Lasky et al., 2012). The proportion of explainable genetic variance attributed to confounded variance correspondingly decreased in the regional models compared to the full range model, although the confounded variance still accounts for the majority of explainable genetic variance. These findings suggest that regional modeling strikes a balance between over- and under-correcting for population structure when parsing how environmental factors pattern genetic variation over complex landscapes. Thus, performing GEA at multiple spatial scales is a valuable strategy that will be especially relevant to similar resequencing studies for many widespread taxa that exhibit comparably strong population structure aligned with environmental variation.

### Major axes of climate variation differ across regions

Through both PCA and RDA approaches, we find that different climate parameters explain the most environmental variation among populations over the full range and in each of the three regional datasets. Moreover, the parameters associated with the most and most significant LFMM hits vary among these groupings. Our capacity to reveal these differences emerges in part because we augmented the often used BioClim variable set with additional variables obtained from the Basin Characterization Model (Flint et al., 2021; Thorne et al., 2015). This model estimates hydrological system inputs and outputs in a manner that allows for more consistent parameter comparisons across habitats with diverse soil conditions and water regimes.

Annual *M. guttatus* populations illustrate the value of this approach because their habitats vary greatly in the form and seasonal timing of available water (FitzPatrick et al., 2023; Kooyers et al., 2015). While some populations rely upon rainfall and associated runoff as their primary water source, others rely primarily on snowmelt from an accumulated winter snowpack, and these differences vary regionally. For example, the OR regional RDA model and LFMM identify Group4 (“Snow”) and Growing Season Length, respectively, as the most prominent agents of climate selection. In contrast, the NC and SC regional RDA models point to Group2 (“Temp in Rainy Season”) and Elevation, respectively, and the NC and SC regional LFMMs identify Group6 (“Precipitation”) and Soil Moisture, respectively, as the primary likely selective factors. Together, we conclude that inclusion of both BioClim and BCM models may improve the ability to discern specific climate components that most vary among populations and that explain genome-wide or locus-specific patterns of genetic variation. However, as both these data sources average climate values over spatial scales generally greater than the typical area covered by a population of *M. guttatus* or other herbaceous annuals, further resolution gains and sensor-based calibrations may yet improve these environmental data sources.

### Substantial incongruence in targets of selection across regions despite sufficient segregating variation

Because we found that climate factors vary substantially in their importance as agents of selection across the range, we predicted that we would find more incongruence than congruence in the genetic targets of climatic selection at a regional scale. Both multi- and univariate GEA analyses of full-range and regional models yielded results consistent with this hypothesis (Figure 4). These results indicate that requiring genotype-environment associations to be detected in multiple population samples can be over-conservative depending on the aims of a GEA study. While this congruence test may yield high-quality candidate genes for downstream investigations of adaptation to one or few shared environmental axes, as seen for the limited set of shared GEA hits in our dataset, it will exclude true positives for region-unique locally adaptive variants related to those or other important factors (Forester et al., 2016; Lasky et al., 2023; Lotterhos, 2023). Like in the case of *TAF5*, range-wide analysis may also not prove as informative about the specific agents of selection related to GEA hits as regional analyses. Moreover, the limited overlap between hits detected by our FR and regional models highlights the value of applying sampling and analysis approaches that mitigate the impact of strong population structure that covaries with the regional importance of different climatic agents of selection. Solely including full range GEA hits when estimating genomic offset for predictive modeling like genomic forecasting, or when designing agricultural or conservation interventions like assisted gene flow, will inevitably then exclude many informative markers that contribute to fitness or yield at the regional scale (Ahrens et al., 2026; Breithaupt et al., 2026; Fitzpatrick et al., 2026; Grabowski et al., 2025; Pegard et al., 2026; Rumberger et al., 2026).

At the level of an adaptive SNP or haplotype, failure to observe a shared genotype-environment association across multiple regions could result from several biological or methodological reasons, but the relative contributions of these sources are rarely evaluated. One possibility is that a variant adaptive in one region is absent or fixed in another region, or at least present at too low or too high a frequency due to sampling, recent introduction by gene flow, young allele age, or other causes that an association cannot be evaluated (Duan et al., 2026). When we examined the allele frequency spectra of variants that did not have significant genotype-environment associations by in a focal pRDA model (FR, OR, NC, or SC) but did in one or more of the other models, we found that around 60-90% of variants segregated at intermediate frequencies at which a signal was potentially detectable (Figure 4c-f; variation in detectability thresholds and proportions likely relates to the size and structure of regional sampling, Figures S22-S23).

Thus, while some SNP-level incongruence was due to the lack of sufficient segregating variation, the majority of the incongruence observed instead appears to result from a lack of selection or response to selection. These examples of incongruence may result from absence of parallelly structured agents of climatic selection in other regions; local adaptation to a similar selection gradient but a nonmonotonic relationship between allele frequency and the environment; the presence of a similar selection gradient but a response involving other loci due to selection on alternate substitutable or higher effect-size alleles; lack of allelic expression in regionally common genetic backgrounds; or stronger regional collinearity of population structure and environment, among other causes (Bishop et al., 2025; Booker, 2024; Duan et al., 2026; Lasky et al., 2023). Some simulation studies have evaluated the relative impacts of these factors (Bishop et al., 2025; Booker, 2024; Forester et al., 2016; Lotterhos, 2023). However, future experimental studies that are designed to discriminate among these possibilities would be valuable.

### Regulators of the FLC/MAF gene family harbor ample climate and phenotype-associated variation

Simulation studies have suggested that GEA approaches may have difficulty detecting true positive loci related to highly polygenic or pleiotropic genetic architectures, particularly in the context of complex demographic histories like that of *M. guttatus* (Lotterhos 2023).

Nonetheless, we detected 17 shared signals between the GEA and GWA datasets for the modest number of clinally varying phenotypes measured on the individuals newly resequenced as part of this study. Given the genes present in the GEA results, we hypothesize that additional GWA studies of phenotypes known to vary clinally across annual *M. guttatus* populations (e.g., chemical defenses and stomatal density) and other important ecophysiological traits may yield additional shared hits (Kooyers et al., 2015, 2017).

Notably, both the GEA and GWA analyses identified signals of allele frequency associations to climate and phenotype, respectively, for multiple genes homologous to regulators of the *FLC/MAF* gene family. One positive regulator, *SUF4*, came up as significant in both approaches. Although alleles of *FLC/MAF* genes or one upstream regulator, *FRIGIDA*, have frequently been implicated in flowering time variation and adaptation in the Brassicaceae (Fournier-Level et al., 2022; Irwin et al., 2016; Johanson et al., 2000; Kuittinen et al., 2008; P. Li et al., 2014; Madrid et al., 2021; Salomé et al., 2011; Shindo et al., 2005; N. Wang et al., 2011; Werner et al., 2005; Zhai et al., 2024), they are not commonly associated with flowering time adaptation outside this group, and additional regulators of *FLC/MAF* genes are not commonly associated with adaptation in the Brassicaceae. However, *M. guttatus* has a large lineage-specific expansion of this gene family; reference genomes include as many as 12 paralogs (Preston et al., 2016). *Mimulus guttatus* is an obligate long-day flowering species, and while perennial *M. guttatus* populations obligately require vernalization to flower if initially raised under short day conditions, relatively few annual populations do (Friedman & Willis, 2013; Kooyers et al., 2015). Notably, flowering time QTLs have been found that colocalize with a cluster of *FLC/MAF* paralogs (Berg et al., 2026; Friedman & Willis, 2013), and some of the *FLC/MAF* clade genes do exhibit photoperiod- or vernalization-sensitive expression in one vernalization-responsive annual population (Preston et al., 2016). Given these past findings and our current results, we hypothesize that the expansion and diversification of the molecular expression and functions of this gene family in *M. guttatus* provide considerable opportunity for adaptation through both *cis*- and *trans*-regulatory variants.

## Data Accessibility

The raw sequencing data and processed assemblies used in this work have been deposited in the NCBI Sequence Read Archive (SRA) under BioProject accession PRJNA1529301. All code used in this project can be found at https://github.com/BlackmanLabUCB/MimulusORCC.

## Benefit Sharing

Benefits Generated: Benefits from this research include the sharing of our genome resequencing data and code on public databases as described above. In addition, as the resequencing data were generated in part with funding from the California Conservation Genomics Program, these data will be included in a broader survey of genetic diversity across many Californian taxa to identify areas of conservation concern.

## Author Contributions

RNH, JMC, JHT, NJK, JPS, BKB designed the research. RNH, MS, RMB, JMC, VC-G, NJK, and BKB performed the research. RNH and BKB contributed to data analysis and visualization. RNH, NJK, and BKB wrote the original draft of the manuscript. RNH, MS, JPS, JHT, NJK and BKB contributed to review and editing of the manuscript. RNH, VC-G, JPS, JHT, NJK, and BKB obtained funding for the research. BKB supervised the research and project administration.

## Acknowledgments

We thank L. Wei, C. Robeson, C. Fang, M. Meek, and C. Kortleven for their contributions to plant husbandry, phenotyping, and DNA extraction. We also thank D. Tataru, K. Ferris, A. Gaudinier, and J. Willis for contributions of seed collections, and C. Wistrom for assistance with maintaining our growth facilities. We thank A. Flint and L. Flint for their contributions to the Basin Characterization Model datasets. We also thank D. Runcie, S. Brady, L. Breithaupt, S. Holt, F. Deligne, and members of the Blackman Lab for helpful discussions and expert advice as the project developed, as well as B. Williams, M. Nachman, and J. Vogel for their insightful comments and constructive feedback on early drafts of this manuscript.

This work was supported by grants from the National Science Foundation to BKB (IOS-1558035, IOS-2222464), JHT (IOS-2222465), NJK (DEB-2045643, IOS-2222466), and JPS (IOS-2222467); a National Institutes of Health Ruth L. Kirschstein National Research Service Award Individual Postdoctoral Fellowship to JMC (F32 GM125244-01); an award to BKB and JPS from the California Conservation Genomics Project, with funding provided to the University of California by the State of California, State Budget Act of 2019 [UC Award ID RSI-19-690224]; and a Student Research Grant to RNH from the California Native Plant Society. Additional support was provided by the Undergraduate Research Assistantship Program, the Sponsored Program for Undergraduate Research, including a Student-Initiated Project award to VC-G, and a Rose Hills Summer Scholarship to VC-G from the SURF L&S Program at the University of California, Berkeley.

## Supplementary Information

### Supplemental Methods

#### Sample collection

We obtained re-sequencing data for 327 individuals from 110 annual populations of *Mimulus guttatus* sampled across Oregon and California [Table S1], 237 of which were sequenced as part of the California Conservation Genomics Project (Shaffer et al., 2022). Sequences for the remaining 90 individuals can be found in the NCBI SRA database under the BioProject PRJNA776633 (Colicchio et al., 2021). The 237 newly re-sequenced individuals in our sample set were grown mostly from existing or newly gathered field-collected seed but were in some instances sampled from later generations [Table S1]. These individuals were grown in a walk-in growth room set to 16 h, 21 °C days: 8 h, 18 °C nights and 50% humidity (Environmental Growth Chambers).

#### Sample preparation and whole-genome resequencing

For the 237 newly re-sequenced samples, DNA was extracted from frozen bud tissue stored at -80°C following a modified CTAB protocol (Kelly & Willis, 1998), quantified using Quant-iT™ PicoGreen™ dsDNA Assay Kits (ThermoFisher), and diluted to ∼2.5 ng/μl.

Individually barcoded libraries were created for each sample using the plexWell™ LP384 Library Preparation Kit (SeqWell). We confirmed library fragment size distributions by Bioanalyser and quantified the purified libraries with the Qubit dsDNA HS Assay Kit (ThermoFisher). Libraries were sequenced 150 bp PE reads to an average depth of ∼15x coverage using the Illumina NovaSeq 6000 instrument available through the California Institute for Quantitative Biosciences at UC Berkeley QB3 Genomics Core (RRID:SCR_022170).

#### Read mapping and variant calling

Reads were trimmed and aligned using the Paleomix pipeline (Schubert et al., 2014). Trimming was performed with the AdapterRemoval option (Lindgreen, 2012), and alignment was performed using BWA-mem (H. Li, 2013) to map reads to the *M. guttatus* var. IM62 v3.1 reference genome [https://phytozome-next.jgi.doe.gov/info/Mguttatusvar_IM62_v3_1; (Lovell et al., 2025). Variant calling was performed on resulting bam files following the GATK best practices workflow (Van der Auwera, 2020; Van der Auwera et al., 2013). Briefly, individual GVCFs were first generated with “HaplotypeCaller” (Poplin et al., 2018) and then combined into one cohort GVCF with “CombineGVCF”. For some population genomic analyses, GVCFs including nonvariant sites for select genome statistics were filtered (--min-base-quality-score 25 --max-alternate-alleles 1 -ploidy 2 --heterozygosity 0.02). For all other analyses, a preliminary cohort vcf containing only biallelic SNPs was generated using “GenotypeGVCFs”. Two filtered vcf datasets were created: one using hard filtering and one using the VQSR method.

Hard filtering was performed using the following “VariantFiltration” parameters (QD < 2.0; QUAL < 30.0; SOR > 3.0; FS > 60.0; MQ < 40.0; MQRankSum < -12.5; ReadPosRankSum < - 8.0). SNPs were further filtered with vcftools (Danecek et al., 2011a), retaining only biallelic SNPs with missing data >60% in individuals with <10% missing data.

To create the truth dataset of high-quality SNPs required as input for VQSR, more stringent hard filtering using “Variant Filtration” was performed on the GVCF using the following parameters (QD < 2.0; QUAL < 30.0; ExcessHet < 54.69; SOR > 1.0, MQ < 50.0; -1.0 > BaseQRankSum < 1.0; MQRankSum < -12.5; 0 < DP < 5.0). The truth dataset was used to train a model and generate VQSLOD scores for 70.0, 90.0, 93.0, 95.0, 97.0, 99.0, 100.0 tranches across 1,2,4,6 gaussians. Out of these 28 iterations, we chose the 90.0 tranche using 4 gaussians to maximize the number of true positive SNP calls without excessive false positives. This VQSR-filtered dataset was then used as input for most analyses.

#### Population genomic parameters

Nucleotide diversity (π) was estimated from gvcf files containing both variant and invariant sites for each population using the “–site-pi” command in VCFtools v0.1.12a (Danecek et al., 2011b). Genetic differentiation (F_ST_) was calculated between each pair of the three regional population groups using VCFtools v0.1.12a using the “--weir-fst-pop” parameter. For each of our three ancestry groups, we also calculated Tajima’s D in 1kb fixed windows using the “--TajimaD 1000” parameter in vcftools v0.1.12a. Linkage disequilibrium between SNP pairs was calculated for our full range dataset using “--r2 --ld-window-r2 0 --ld-window 99999” parameters in PLINK version1.9. Additional haplotype length selection metrics, integrated Haplotype Score (iHS) and Extended Haplotype Homozygosity (EHH), were calculated for the full range dataset using the R package rehh (Gautier et al., 2017). Genetic data was input as a vcf using the R package vcfR (Knaus & Grünwald, 2017), and iHS was calculated using data2haplohh(), scan_hh(), and ihh2ihs() commands with default parameters. We then ran calc_candidate_regions using “threshold = 1.3, pval = TRUE, window_size = 1, overlap = 0, min_n_extr_mrk = 5” to extract individual iHS outliers with a p value < 0.05. Taking the original haplohh object output from data2haplohh, EHH was calculated around focal SNPs using calc_ehhs with default parameters.

#### Metrics of growing season timing

Because snowpack influences growing season timing in more northern and alpine populations but not in more southern or lower elevation populations, we categorized populations into “colder” and “warmer” groups with differing metrics for the growing season. For “colder” populations, we defined the growing season start date as the month after January when the average temperature exceeds 4 °C and the slope between months for AET exceeds 0.2. We chose these metrics because 4 °C is the accepted temperature for stratification in growth chamber conditions (Luo & He, 2020; Xu & Chong, 2018), and a slope of 0.2 is reflective of population establishment. In contrast, for the “warmer” populations that never experience temperatures below 4 °C, we defined the growing start date as the month, January or later, after which average temperature exceeds 10 °C and the slope between months for AET exceeds 0.2. Since the growing season ends in a terminal drought for all annual populations, we estimate growing season end date as the month after the slope of monthly AET becomes negative. Growing season length was calculated as the difference in number of months between start date and end date.

#### Genome-Wide Association (GWA) analysis

Germination time was measured as the day on which the majority of the ∼10 planted seeds from the same line per pot had sprouted. Time to flower was measured as the number of days from germination to the first day the first flower opened. On a plant’s day of first flower, we also took the following morphological measurements: plant height; blade length, blade width, and petiole length of first true leaf; node of first flower in relation to the node of the first pair of true leaves; peduncle length of first flower; and total nodes. We also qualitatively scored trichome density of the leaves subtending the node of the first flower on a scale of 0-2 (0 = trichomes, 1 = few trichomes, and 2 = many trichomes). As this is a subjective measure, all trichome scores were scored twice, once by each observer, and then cross-validated. Because phenotypes were not collected for the previously re-sequenced Oregon samples and the sample sizes of the regional data are underpowered for GWA analysis, we only performed the GWA analysis for the full set of 237 individuals. We used the same simple imputation method as our ALGATR analyses and included a kinship matrix constructed with PLINK version1.9 as a covariate to control for population structure. We performed GWAS for each of the 10 phenotypes in GEMMA (Zhou & Stephens, 2012) using both linear mixed models and Bayesian Sparse Linear Mixed Models (BSLMMs) (Zhou et al., 2013). LMMs are standard statistical models used to analyze data with both fixed (population-level) and random (individual-level) effects, and BSLMMs are Bayesian-informed models which explicitly incorporate a “sparsity” assumption, where the model tests whether a small subset of genetic variants that have detectable effects on the phenotype above effects above a background a polygenic architecture of many loci of small effect. Each BSLMM was run four times with different seed values and the following parameters “-bslmm 1 -w 5000000 -s 25000000 -rpace 1000 -wpace 10000”. To be considered an outlier, LMM SNPs must exceed a Bonferroni-corrected Wald test p=0.05 and BLSMM SNPs must have an average posterior inclusion probability (PIP) > 0.01 over the four runs. The reported SNPs were also filtered to exclude sites located within intervals flagged as likely multicopy regions by Paramask (Tjeng et al., 2025).

### Supplemental Figures

**Figure S1:**
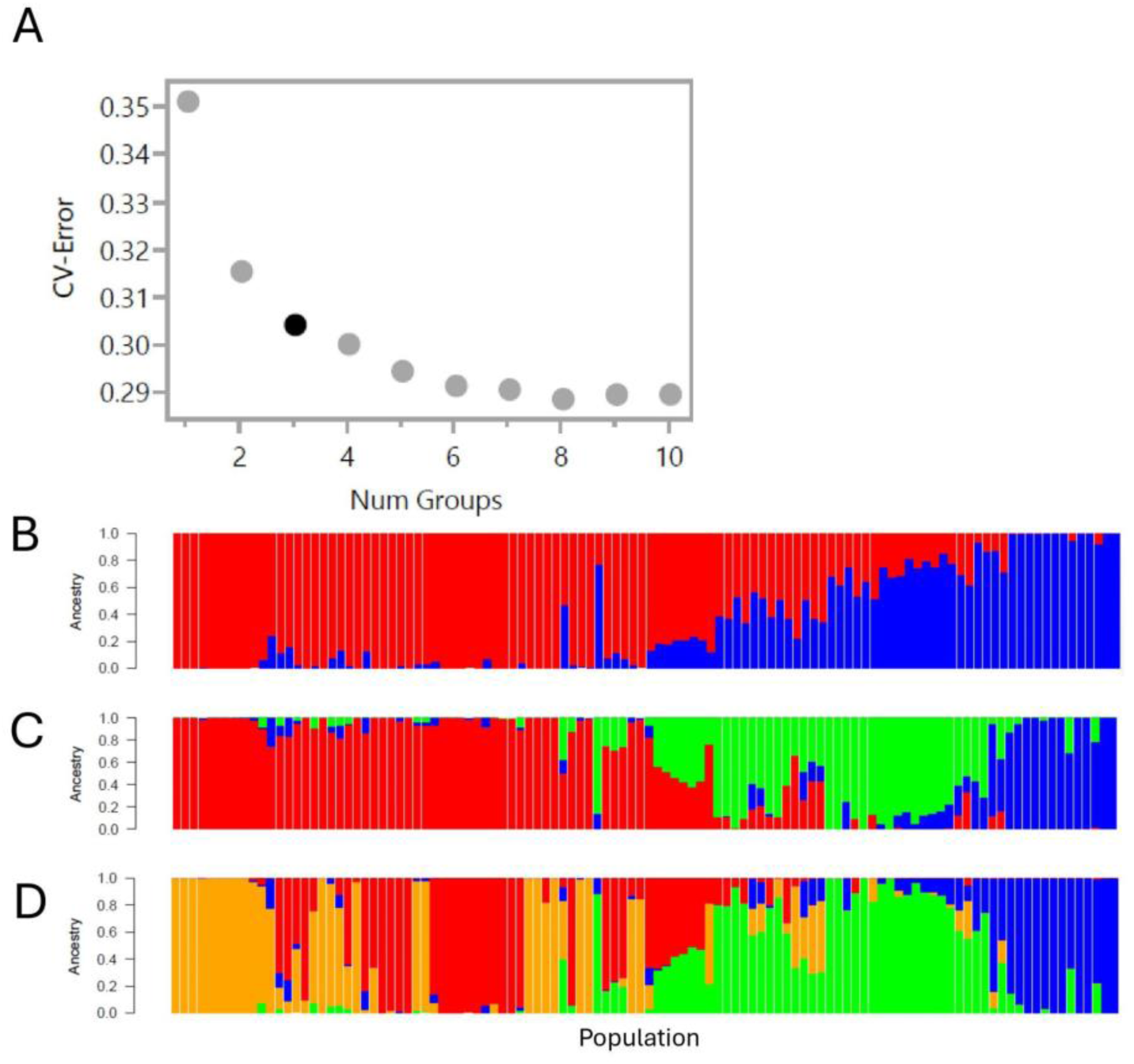
Admixture cross validation error predicts three ancestry groups. (A) Cross Validation Error used to determine the most likely number of predicted ancestry groups (K=3) (B-D) ADMIXTURE population site ancestry arranged by latitude ascending from left to right and colored by ADMIXTURE ancestry grouping for (B) K =2, (C) K = 3, and (D) K = 4.

**Figure S2:**
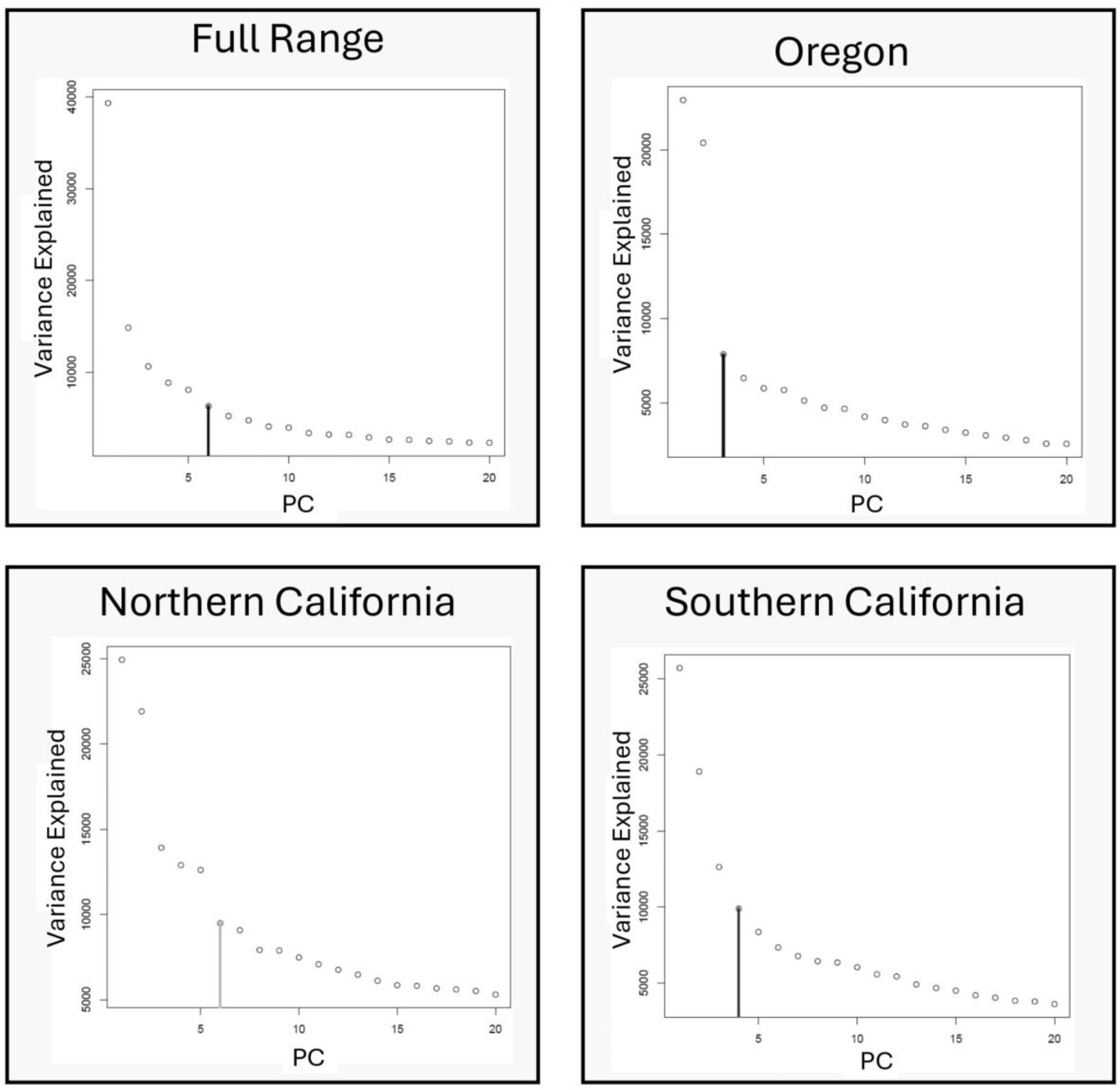
Latent factor estimation elbow plots. Latent Factor Prediction for LFMM model training for each regional Model. Full Range latent factor prediction suggests 6 factors, Oregon latent factor prediction suggests 3 factors, Northern California latent factor prediction suggests 6 factors, and Central and Southern California latent factor prediction suggests 4 factors.

**Figure S3:**
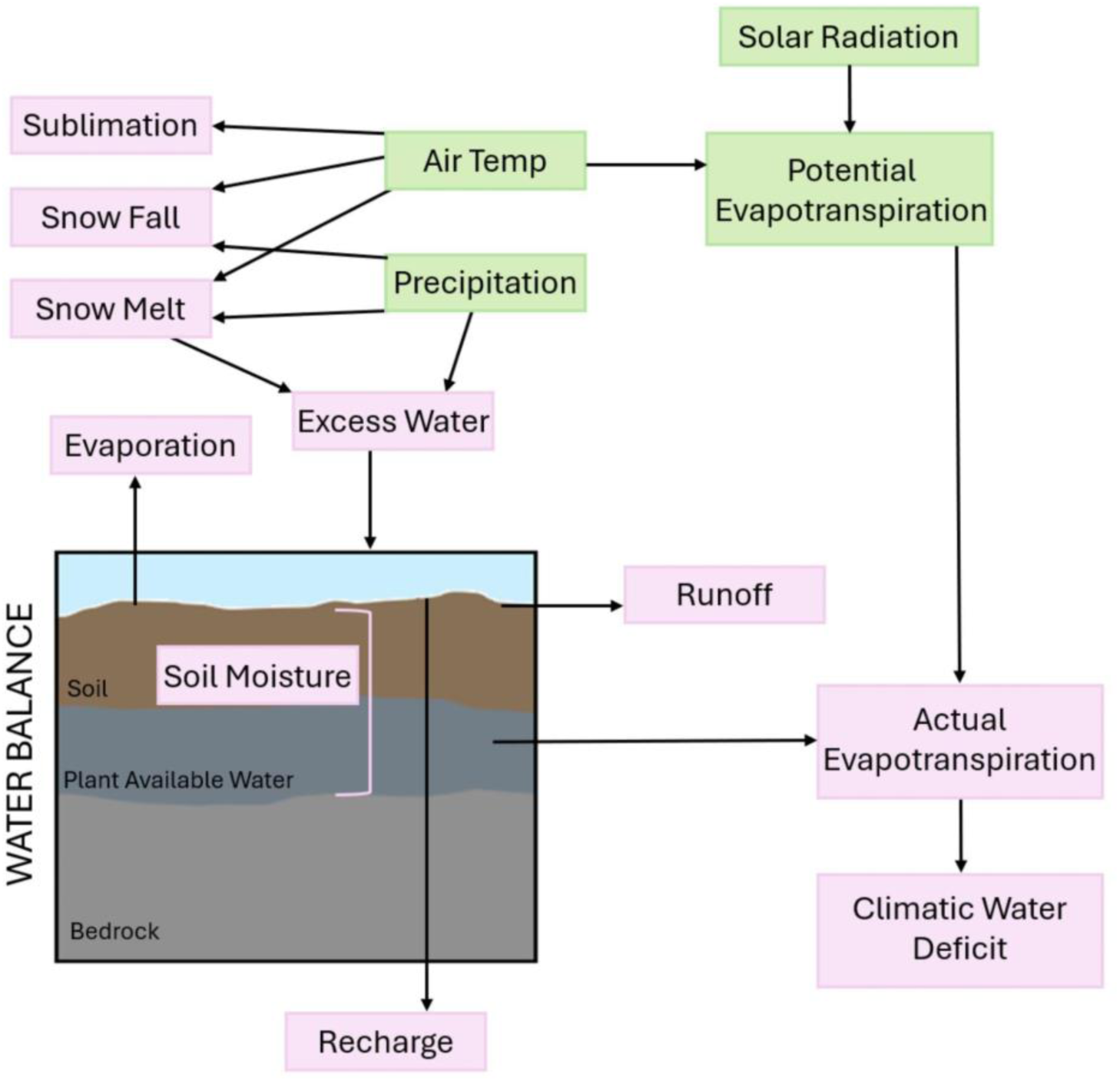
Basin Characterization Model structure. Diagram describing the relationships between Basin Characterization Model environmental variables.

**Figure S4:**
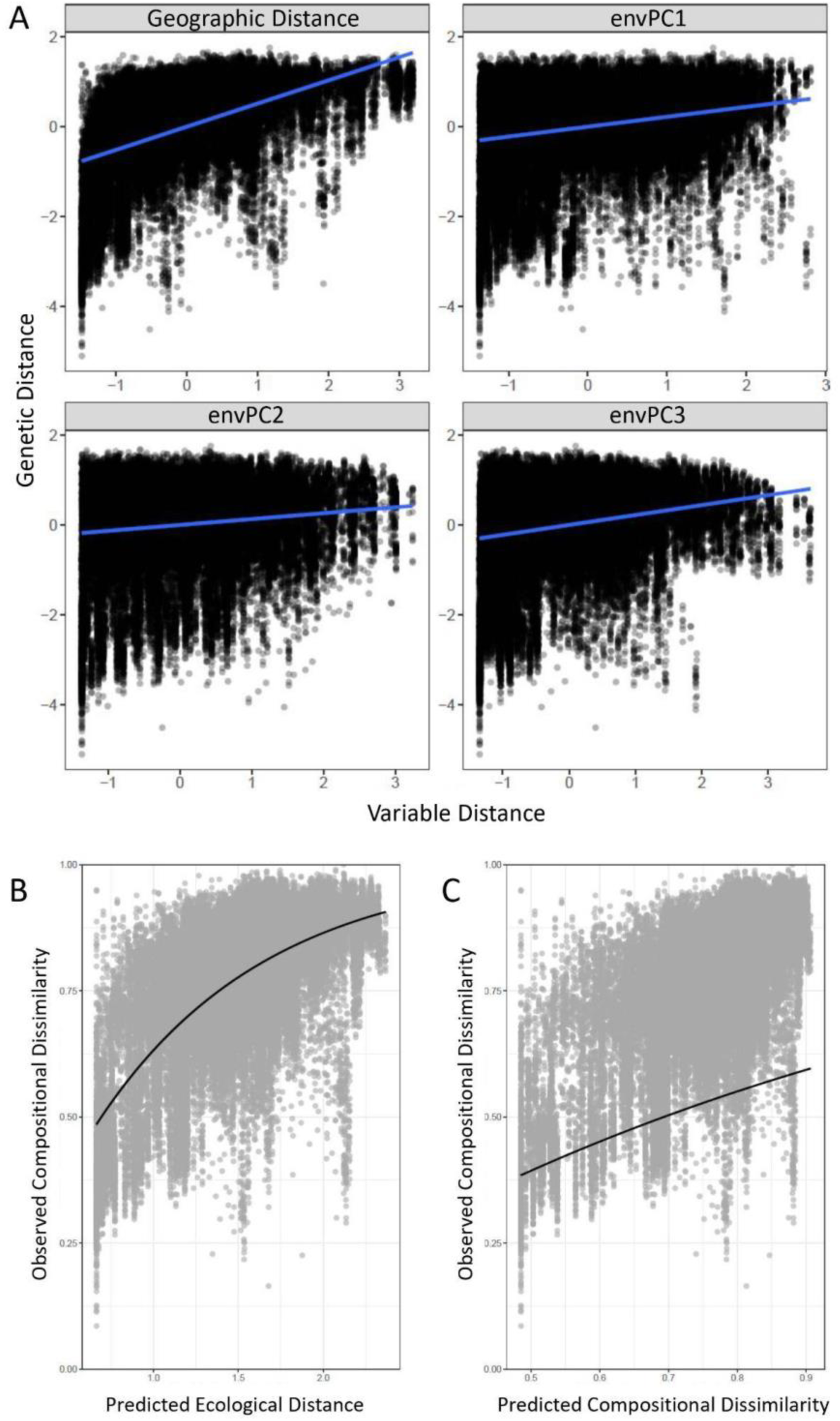
IBD and IBE influence genetic variance across the range. (A) Single variable correlations between genetic distance and geographic distance (a proxy for Isolation by Distance) and the first three environmental variable axes of variation (a proxy for Isolation by Environment) from Full Range Multiple Matrix Regression with Randomization (MMRR). (B) Correlation between observed compositional dissimilarity, defined here as differences in population composition between pairs of locations, and predicted ecological distance, defined here as the raw distance between raw environmental predictor values, from generalized dissimilarity modeling (GDM). (C) Correlation between observed compositional dissimilarity, defined here as differences in population composition between pairs of locations, and predicted compositional, defined here as predicted dissimilarity between population pairs as predicted from environmental dissimilarity, from generalized dissimilarity modeling (GDM).

**Figure S5:**
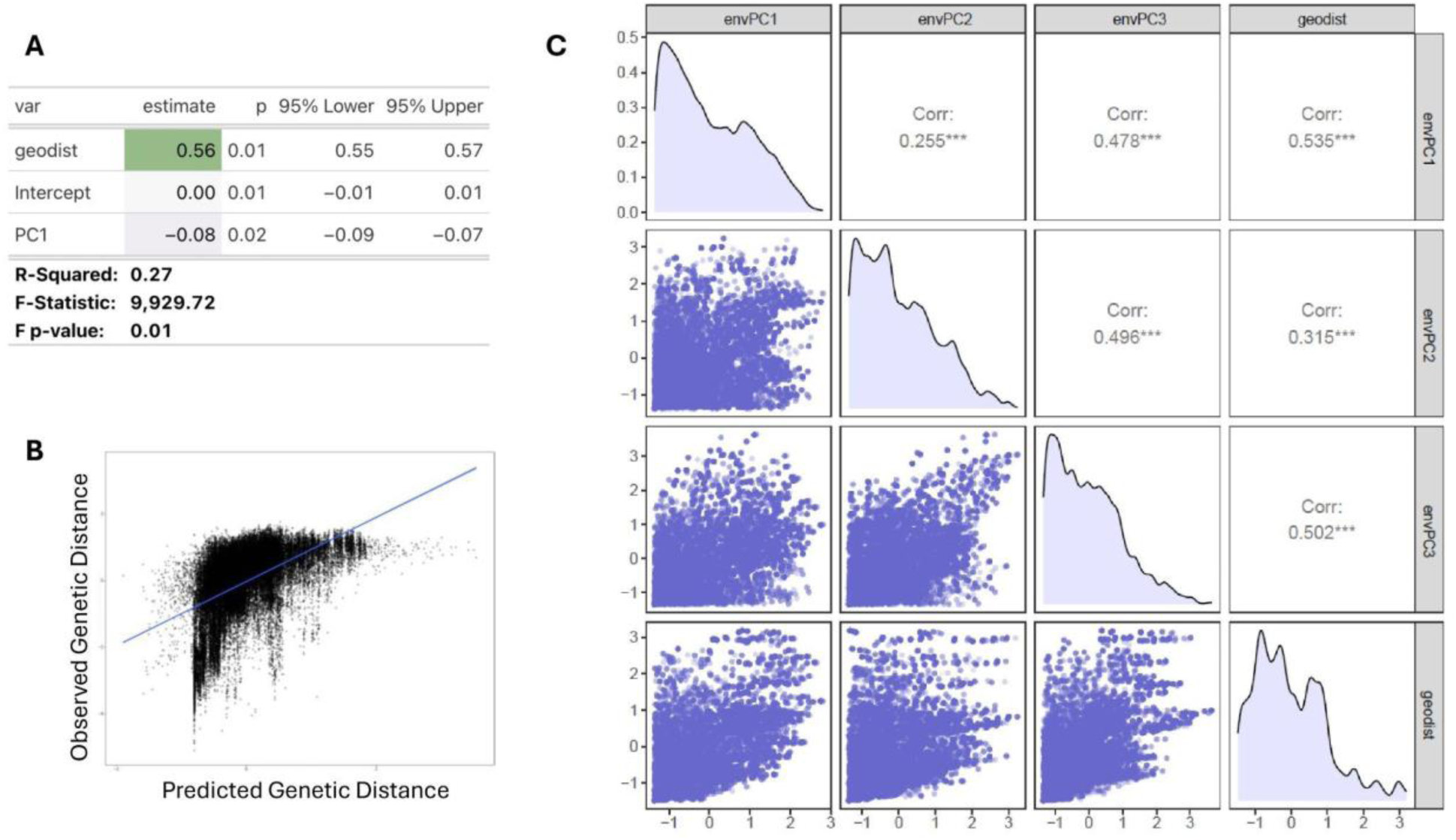
MMRR additional metrics. (A) Multiple Matrix Regression with Randomization (MMRR) coefficients of the variables. Geodist is geographic distance using population gps location. (B) Multiple Matrix Regression with Randomization (MMRR) fitted relationship that compares predicted genetic distance against the observed genetic distance. (C) Multiple Matrix Regression with Randomization (MMRR) covariances between the predictor variables (the first three axes of environmental variance: EnvPC1, EnvPC2, EnvPC3). All three EnvPCs are correlated with geographic distance.

**Figure S6:**
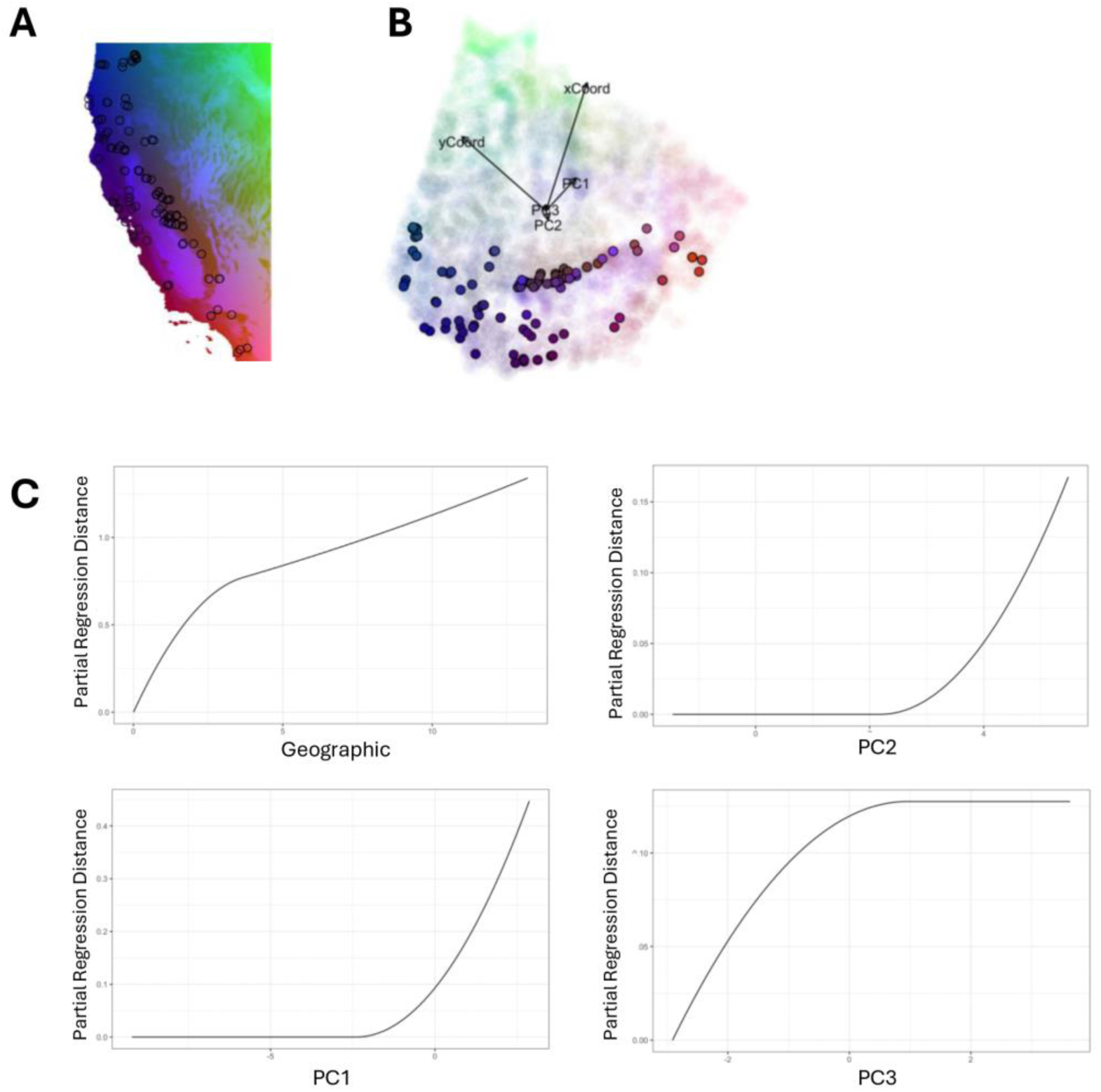
GDM additional metrics. (A) Environmental Variance (EnvPC1, EnvPC2, and EnvPC3) rbg BioClim variable raster overlay. Sampled population sites are denoted by black circles. (B) Generalized Dissimilarity Model (GDM) biplot. PC1, PC2, and PC3 denote the first three axes of genetic variation. Points are colored by EnvPC1, EnvPC2, and EnvPC3 rbg BioClim variable raster overlay. Solid points represent the grid cells in which our population sites occur. Blurred points represent unsampled, but predicted grid cells. (C) Splines from the GDM model comparing Partial Regression Environmental Distance between sites with the first three axes of genetic variation (PC1, PC2, PC3)

**Figure S7:**
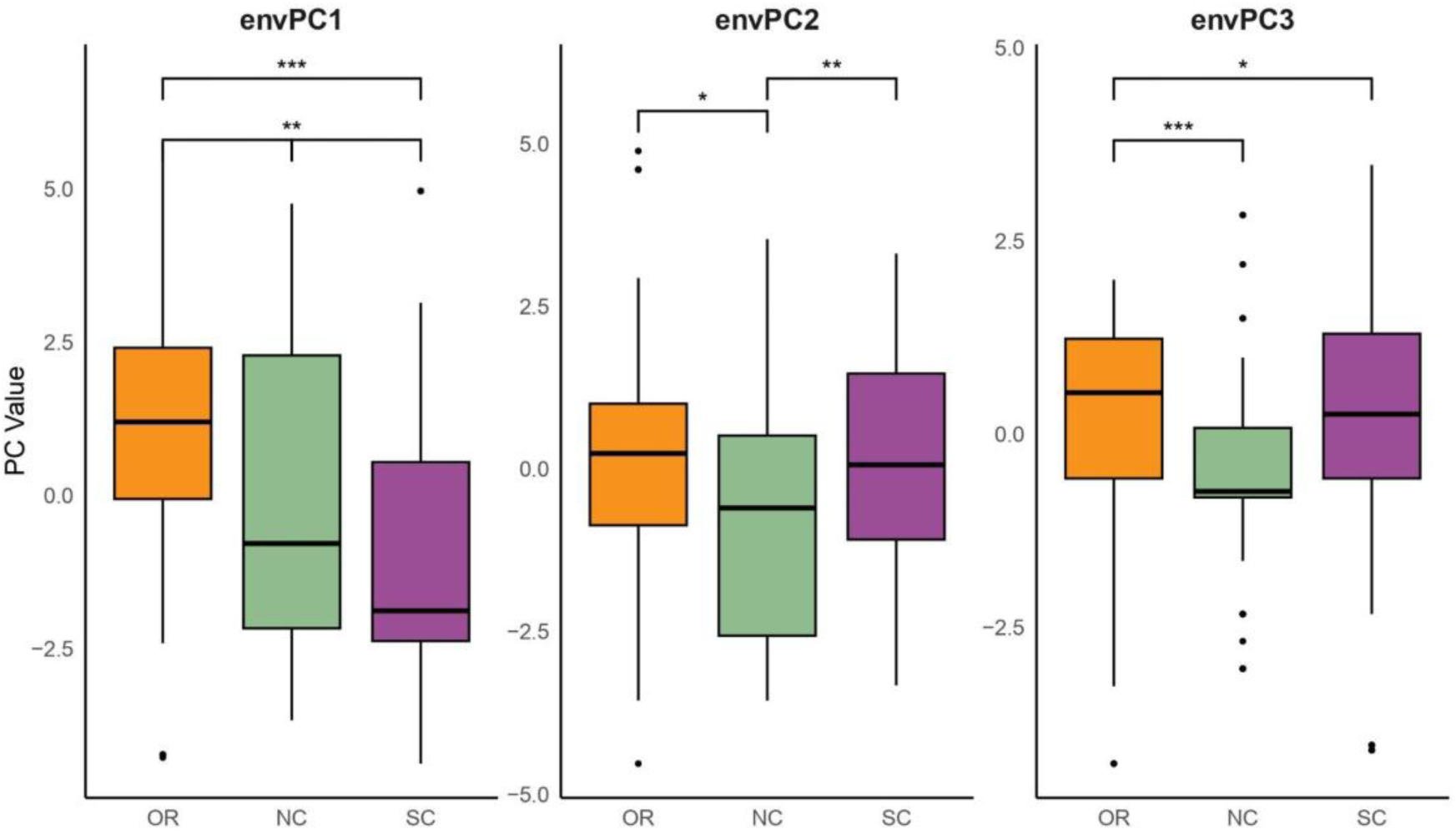
EnvPC1/2/3 t-Tests across regions. (A-C) Pairwise T-test significance across regions for (A) EnvPC1, (B) envPC2, and (C) envPC3. The p value for each comparison is as follows: Oregon and Northern California = 0.0039, Northern California and Central + Southern California = 0.001; Oregon and Central + Southern California = <0.001.

**Figure S8:**
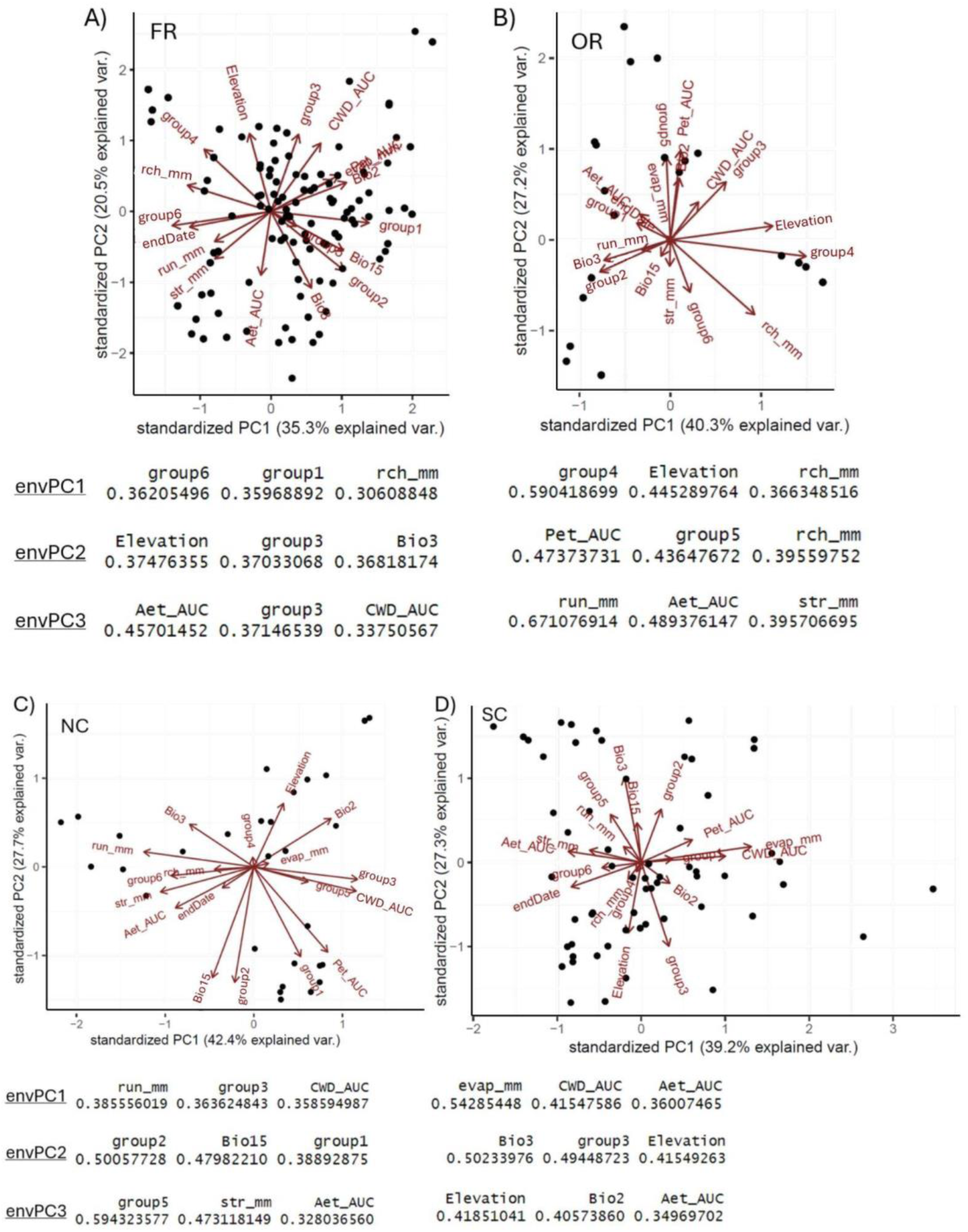
Environmental PCA by regional model. (A) Biplot of environmental variables for the Full Range. Eigenvectors are denoted as red arrows. The absolute value of the EnvPC1 score for the three environmental variables most responsible for EnvPC1 are (Group6 = 0.362; Group1 = 0.360; Recharge = 0.306). The absolute value of the EnvPC2 score for the three environmental variables most responsible for EnvPC2 are (Elevation = 0.375; Group3 = 0.370; Bio3 = 0.368). The absolute value of the EnvPC3 score for the three environmental variables most responsible for EnvPC3 are (Actual Evapotranspiration = 0.457; Group3 = 0.372; Recharge = 0.338). (B) Biplot of environmental variables for Oregon. Eigenvectors are denoted as red arrows. The absolute value of the EnvPC1 score for the three environmental variables most responsible for EnvPC1 are (Group4 = 0.59; Elevation = 0.445; Recharge = 0.366). The absolute value of the EnvPC2 score for the three environmental variables most responsible for EnvPC2 are (Potential Evapotraspiration = 0.474; Group5 = 0.436; Bio3 = 0.396). The absolute value of the EnvPC3 score for the three environmental variables most responsible for EnvPC3 are (Runoff= 0.671; Group3 = 0.489; Recharge = 0.396). (C) Biplot of environmental variables for Northern California. Eigenvectors are denoted as red arrows. The absolute value of the EnvPC1 score for the three environmental variables most responsible for EnvPC1 are (Runoff = 0.386; Group3 = 0.364; Climatic Water Deficit = 0.359). The absolute value of the EnvPC2 score for the three environmental variables most responsible for EnvPC2 are (Group2 = 0.5; Bio15 = 0.48; Group1 = 0.389). The absolute value of the EnvPC3 score for the three environmental variables most responsible for EnvPC3 are (Group5= 0.594; Soil Moisture = 0.473; Actual Evapotranspiration = 0.328). (D) Biplot of environmental variables for Central and Southern California. Eigenvectors are denoted as red arrows. The absolute value of the EnvPC1 score for the three environmental variables most responsible for EnvPC1 are (Evaporation = 0.543; Climatic Water Deficit = 0.415; Actual Evapotranspiration = 0.36). The absolute value of the EnvPC2 score for the three environmental variables most responsible for EnvPC2 are (Bio3 = 0.502; Group3 = 0.494; Elevation = 0.415). The absolute value of the EnvPC3 score for the three environmental variables most responsible for EnvPC3 are (Elevation= 0.419; Bio2 = 0.406; Actual Evapotranspiration = 0.35).

**Figure S9:**
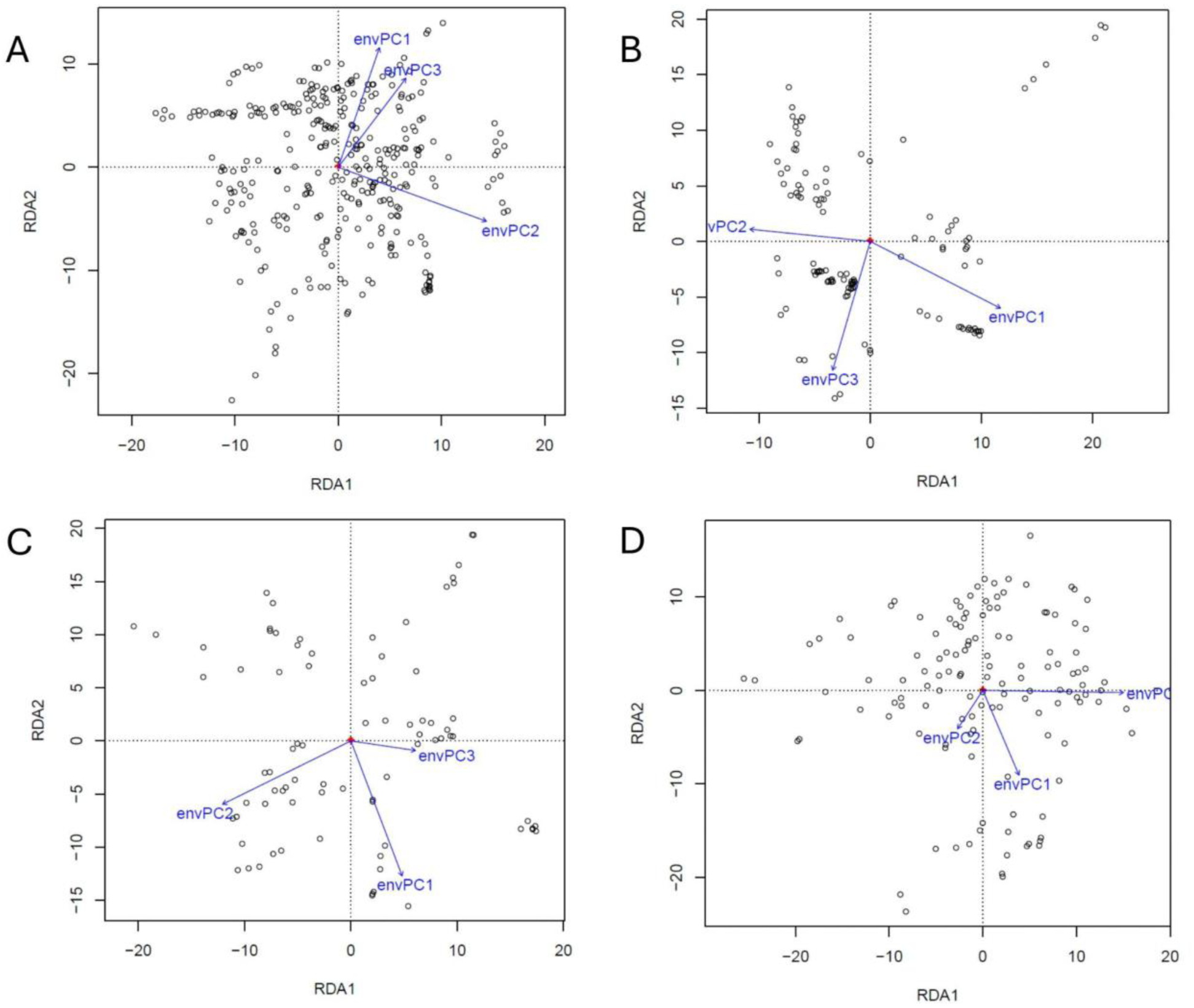
RDA biplots. (A) RDA biplots for the (A) Full Range model, (B) Oregon model, (C), Northern California model, and (D) Central and Southern California model. The eigenvectors for first three axes of environmental variance (EnvPC1, EnvPC2, EnvPC3) are denoted as blue arrows. Individual sample values are denoted as black circles.

**Figure S10:**
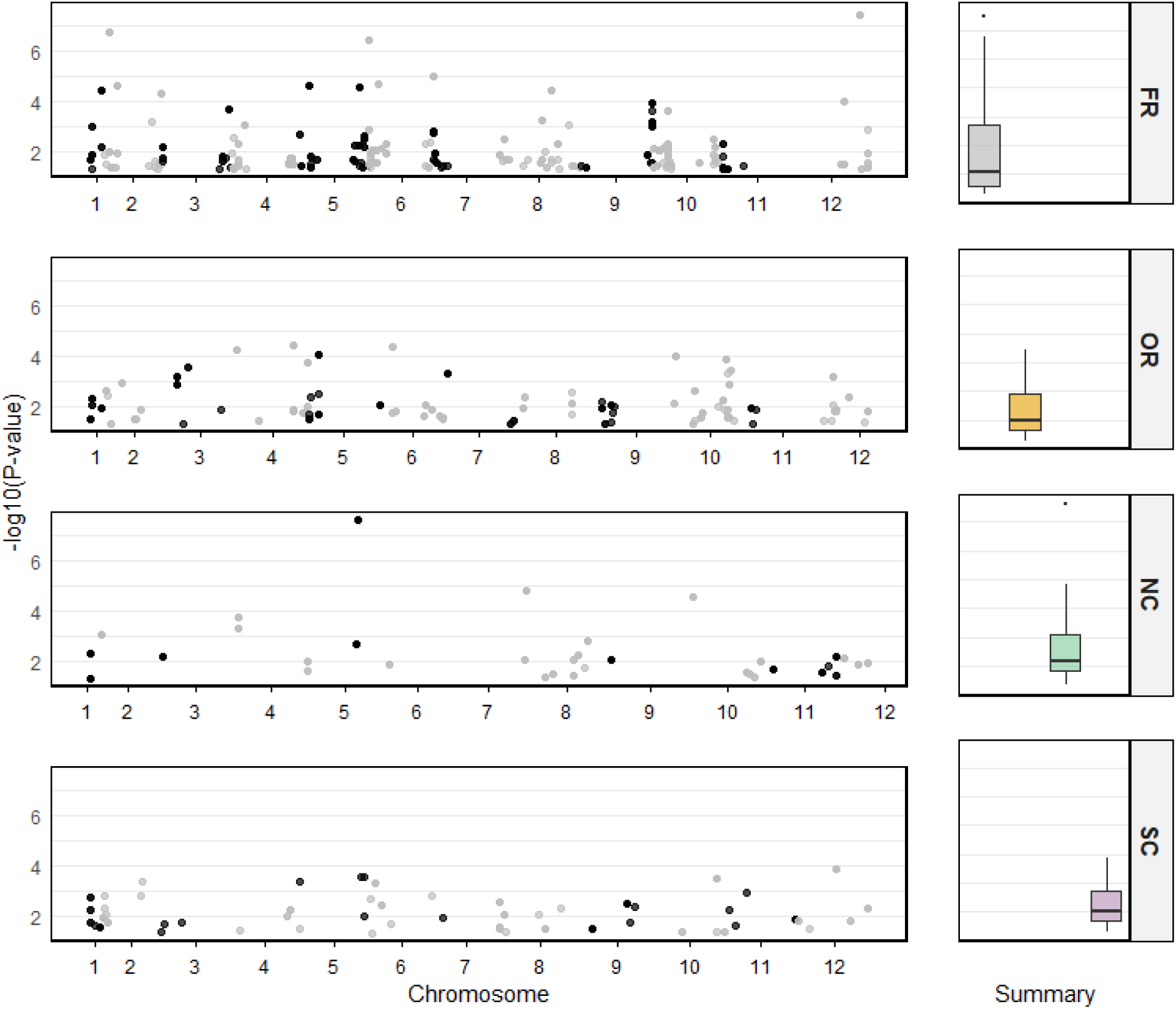
WZA value range for each regional pRDA. Manhattan plots for all significant WZA hits divided by model. Colored box-and-whisker plots indicate value quartiles. Abbreviations: FR = full range, OR = Oregon, NC = Northern California, SC = Central and Southern California.

**Figure S11:**
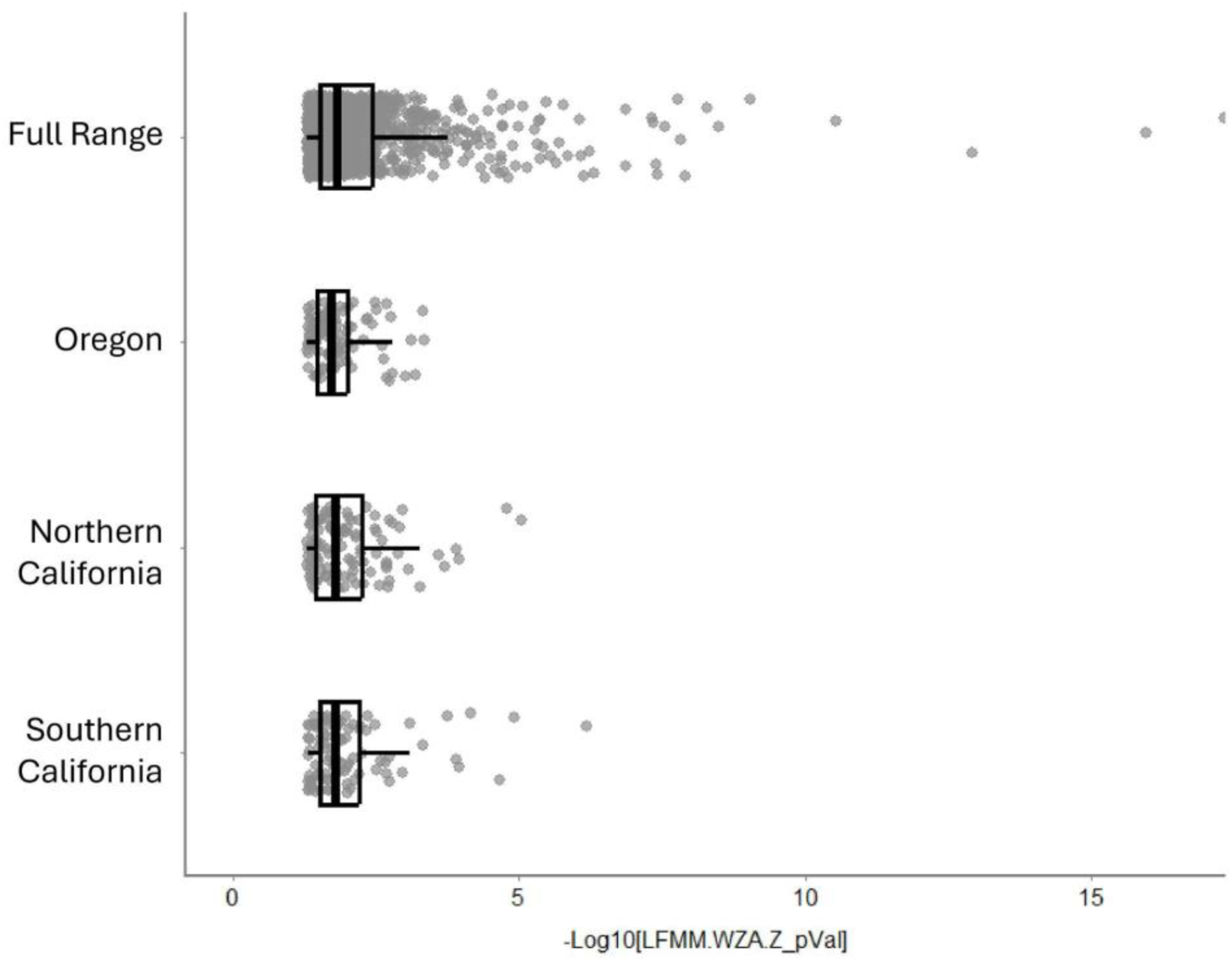
WZA value range for each regional LFMM. Grey dots indicate individual values for each window across each model. Box-and-whisker plots indicate value quartiles.

**Figure S12:**
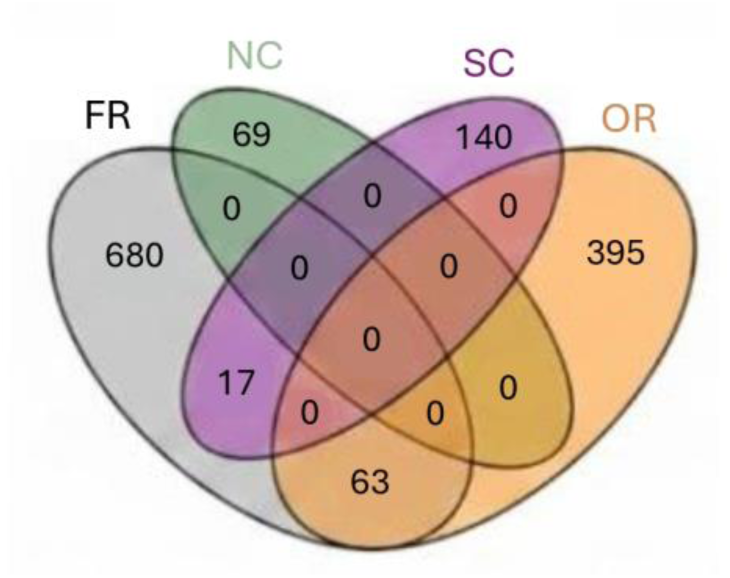
Overlapping SNPs across regional models. Number RDA SNPs which are identified as outliers (p value < 0.05; WZA p value < 0.05) which overlap across RDA regional models.

**Figure S13:**
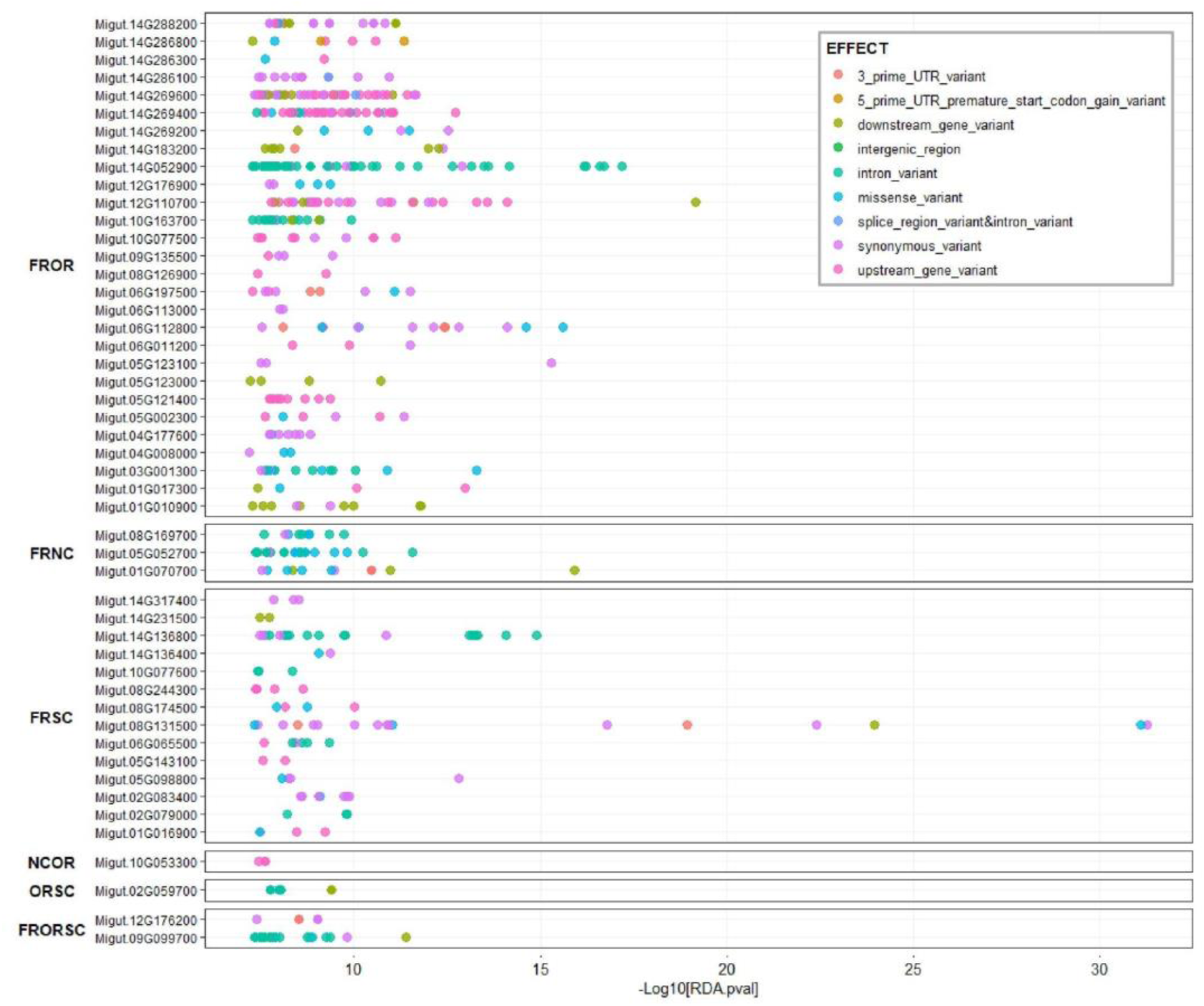
Overlapping genes across RDA regional models. Overlapping genes with significant SNPs across RDA regional models plotted by –log10 RDA p value. Each gene is named using the IM62 v3 annotations. Overlap is denoted by a combination of regional models using the following abbreviations (Full Range = FR, Oregon = OR; Northern California = NC; Central and Southern California = SC). SNPs are colored by the predicted mutation effect.

**Figure S14:**
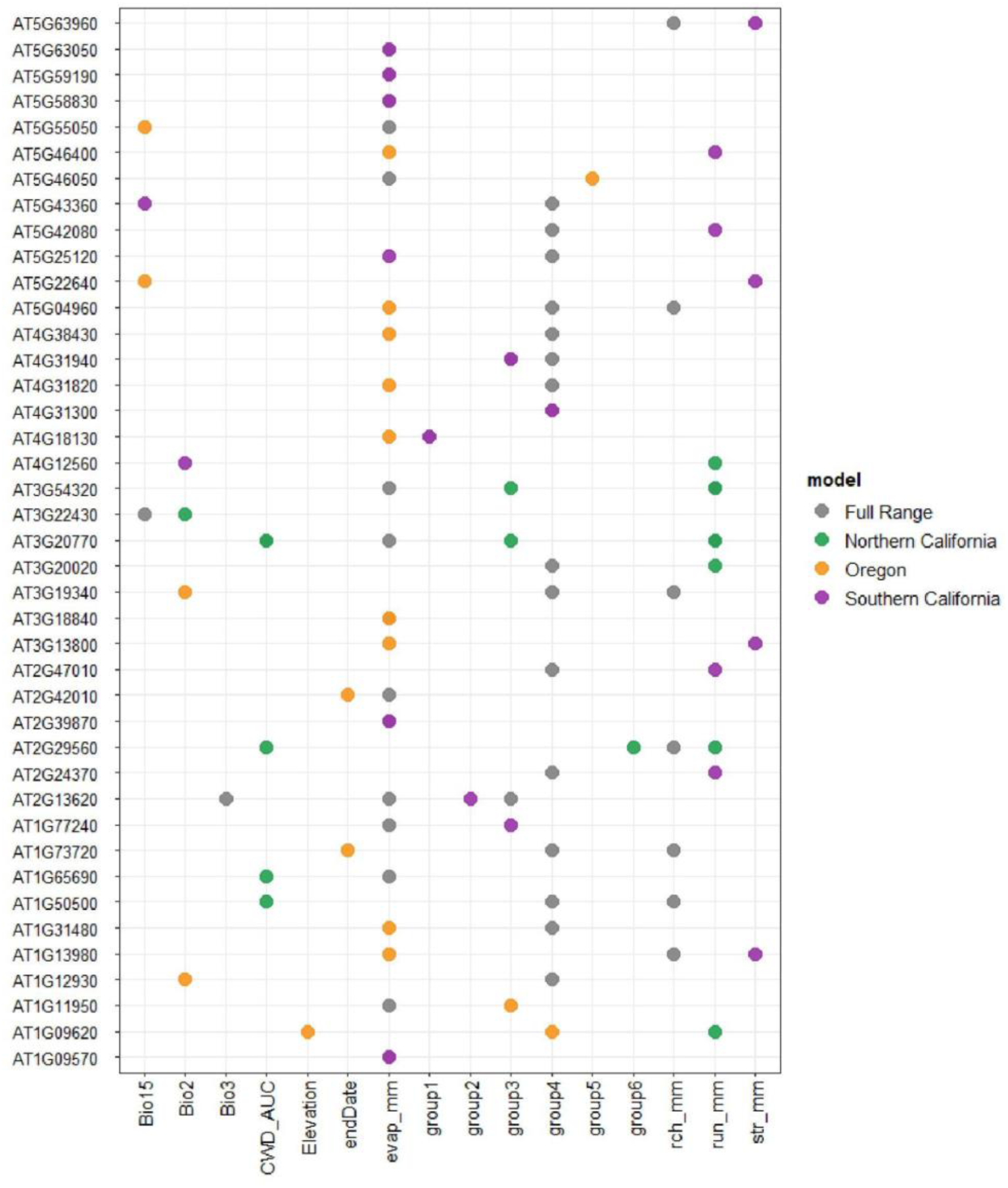
Overlapping genes across LFMM regional models. Overlapping genes with significant SNPs across LFMM regional models plotted by –log10 LFMM adjusted p value and colored by regional model identity. Models. Each gene is named using the *Arabidopsis thaliana* TAIR 10 homolog annotations.

**Figure S15:**
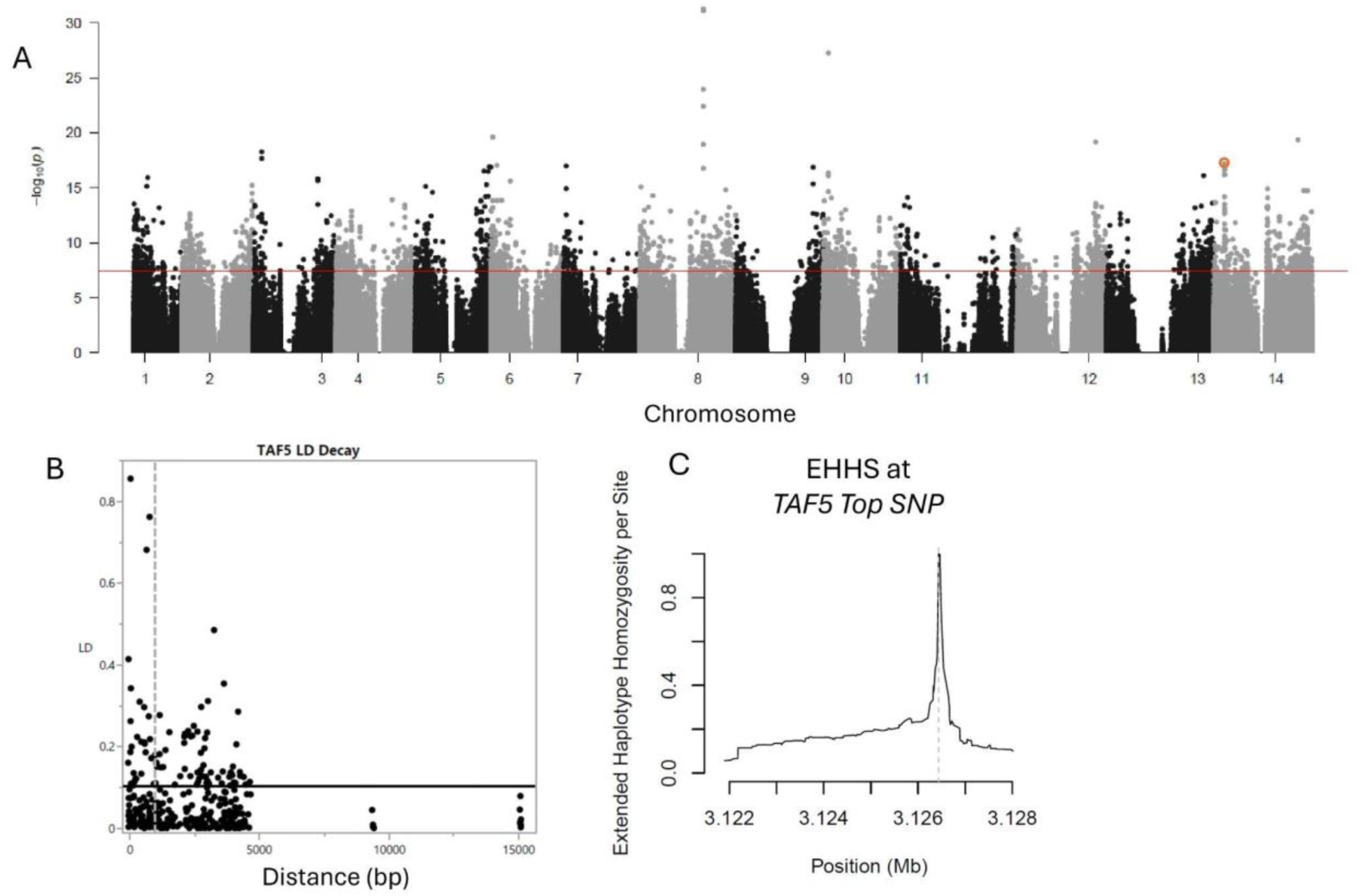
Signatures of Selection on *TAF5*, (A) Whole genome manhattan plot of uncorrected –log10 Full Range RDA p values. The most significant non-synonymous SNP [Chr_14:3126435] is circled in orange. The Bonferroni threshold is represented by the red line. (B) Full Range linkage disequilibrium (LD) decay plotted against the base-pair distance between the most significant non-synonymous SNP and nearby SNPs. Vertical dotted grey line denotes the full genome LD decay estimate. Horizontal black line denotes the threshold defining extended LD. (C) Full Range per site Extended Haplotype Homozygosity (EHH) around the most significant non-synonymous SNP.

**Figure S16:**
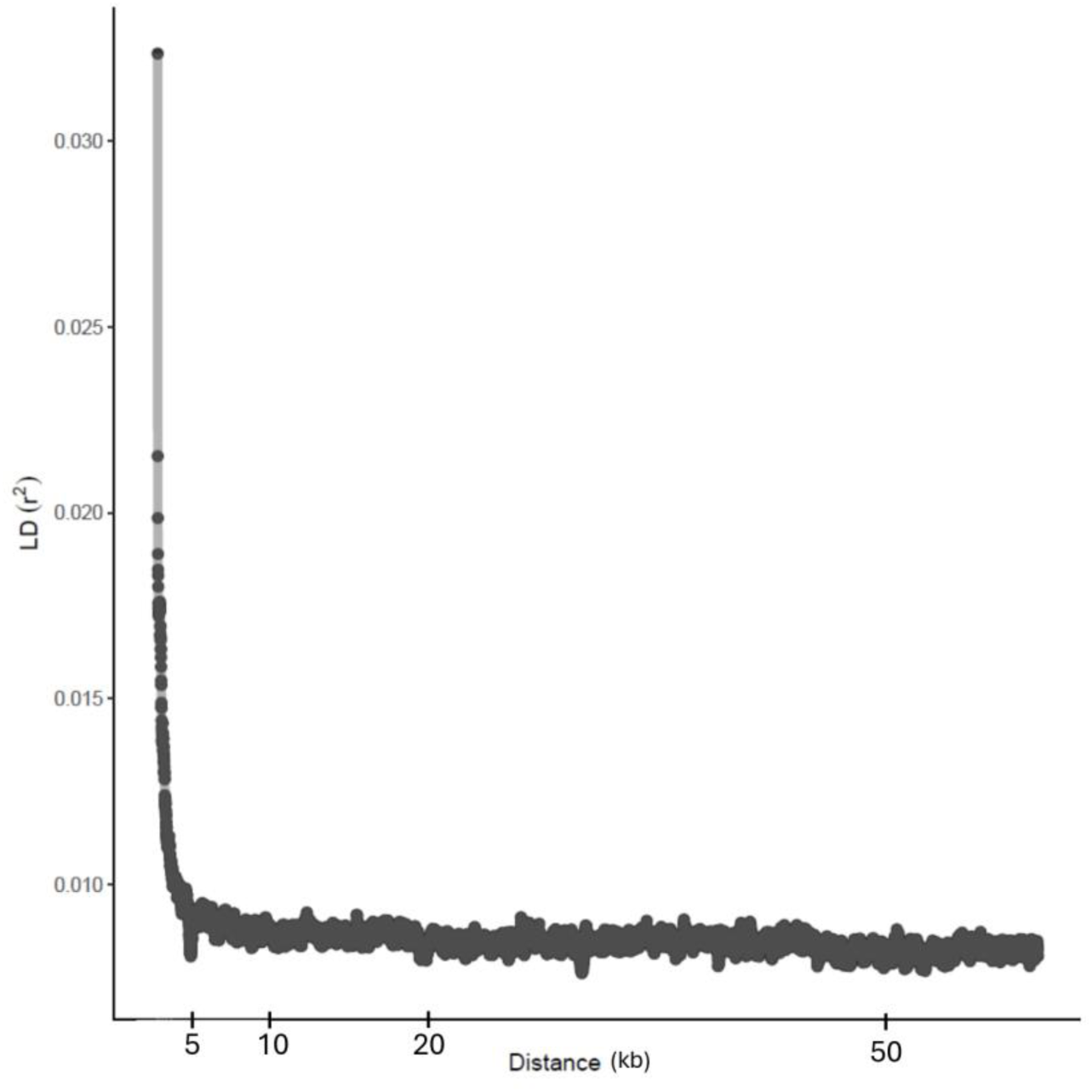
Genomic linkage disequilibrium decay. Linkage disequilibrium (r^2^) plotted against kilobase distance for the whole genome.

**Figure S17:**
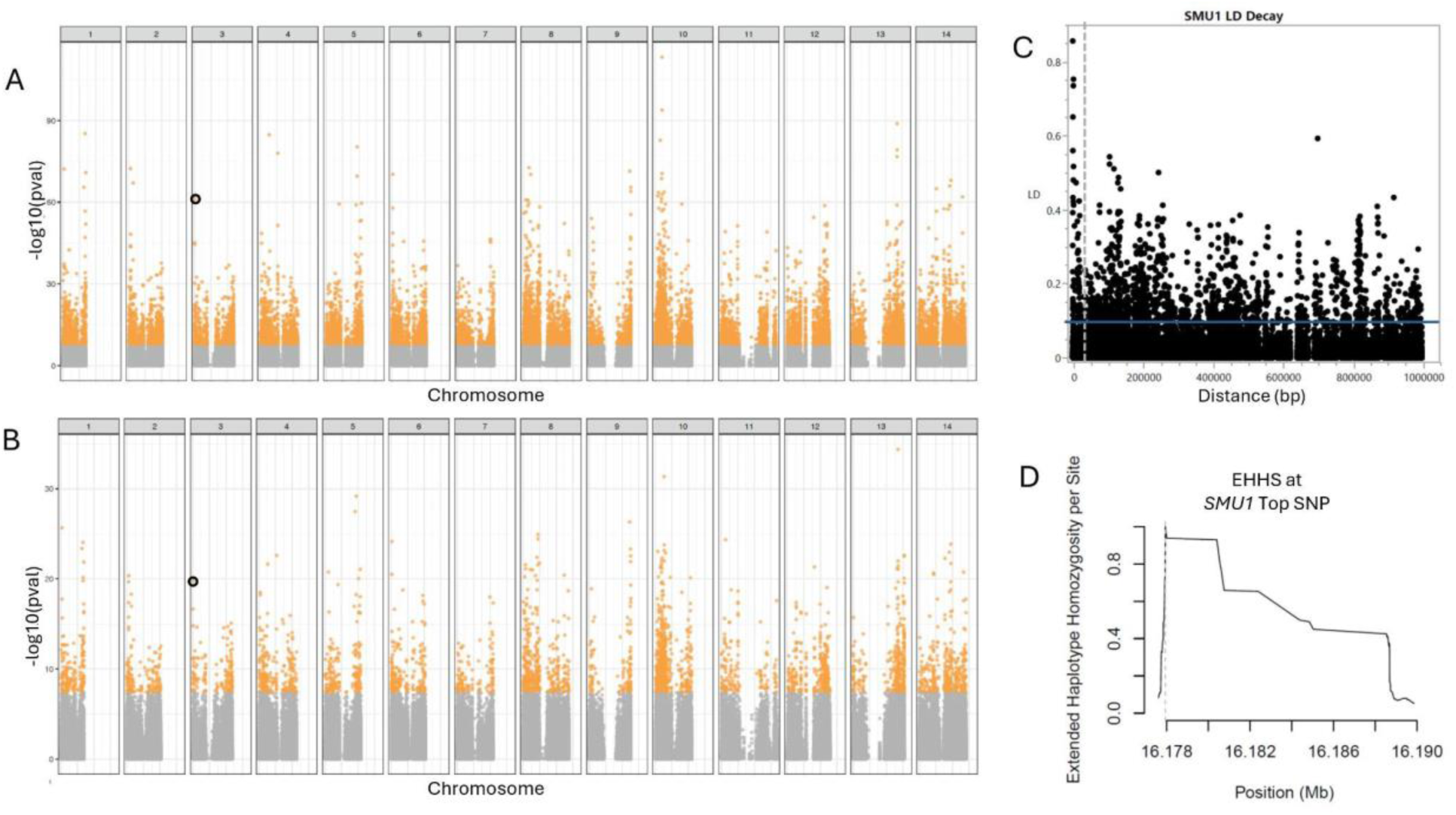
Signatures of aelection on *SMU1*. (A-B) Whole genome manhattan plot of uncorrected –log10 Full Range (A) Group4 (“Snow”) and (B) Recharge LFMM adjusted p values. The most significant non-synonymous SNP [Chr_03:23498841] is circled in black. GEA hits exceeding the bonferroni threshold are colored orange. (C) Full Range linkage disequilibrium (LD) decay plotted against the base-pair distance between the most significant non-synonymous SNP and nearby SNPs. Vertical dotted grey line denotes the full genome LD decay estimate. Horizontal blue line denotes the threshold defining extended LD. (D) Full Range per site Extended Haplotype Homozygosity (EHH) around the most significant non-synonymous SNP.

**Figure S18:**
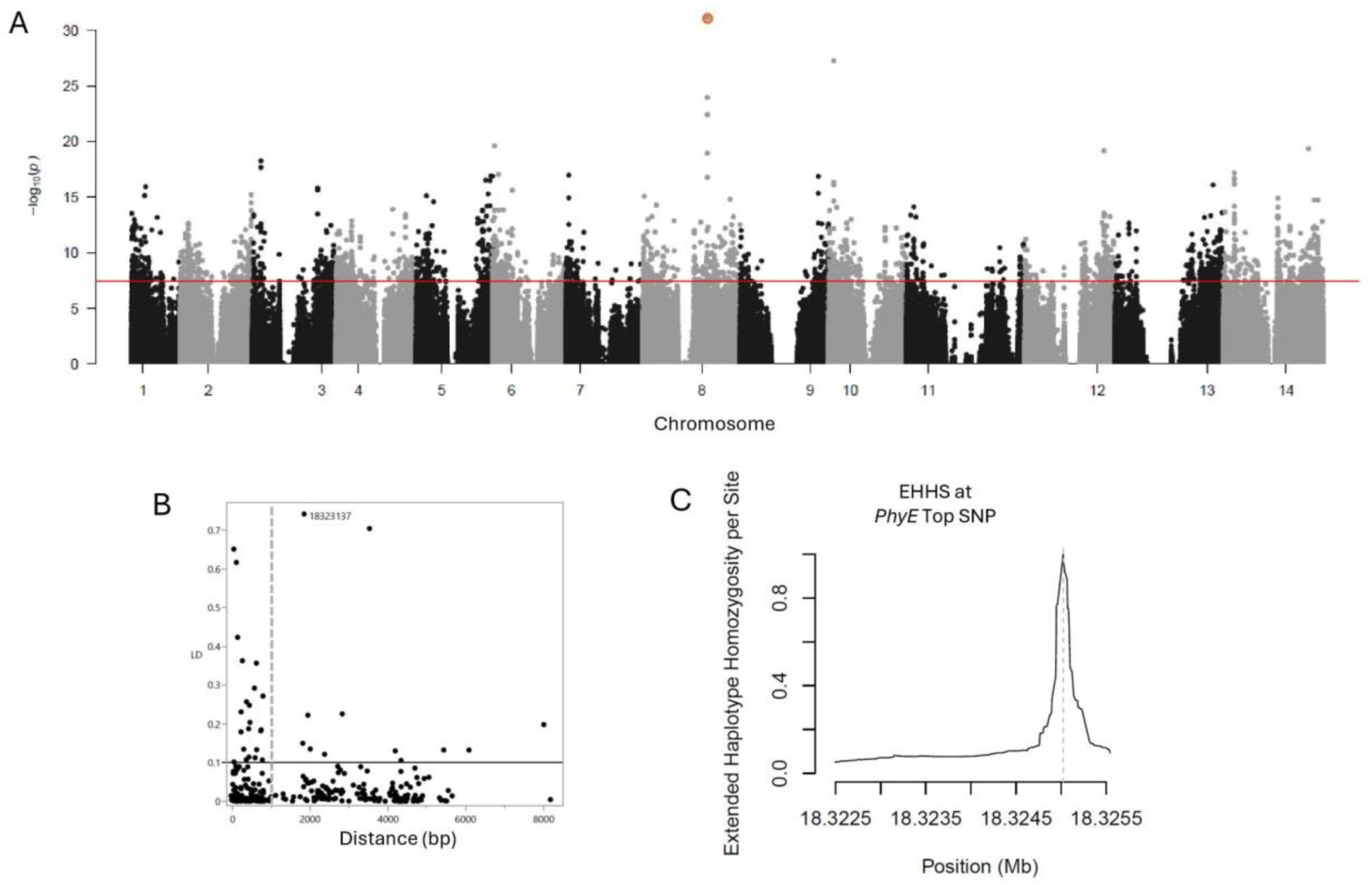
Signatures of aelection on *PhyE*. (A) Whole genome manhattan plot of uncorrected –log10 Full Range pRDA p values. The most significant non-synonymous SNP [Chr_08:18325023] is circled in orange. The Bonferroni threshold is represented by the red line. (B) Full Range linkage disequilibrium (LD) decay plotted against the base-pair distance between the most significant non-synonymous SNP and nearby SNPs. Vertical dotted grey line denotes the full genome LD decay estimate. Horizontal black line denotes the threshold defining extended LD. (C) Full Range per site Extended Haplotype Homozygosity (EHH) around the most significant non-synonymous SNP.

**Figure S19:**
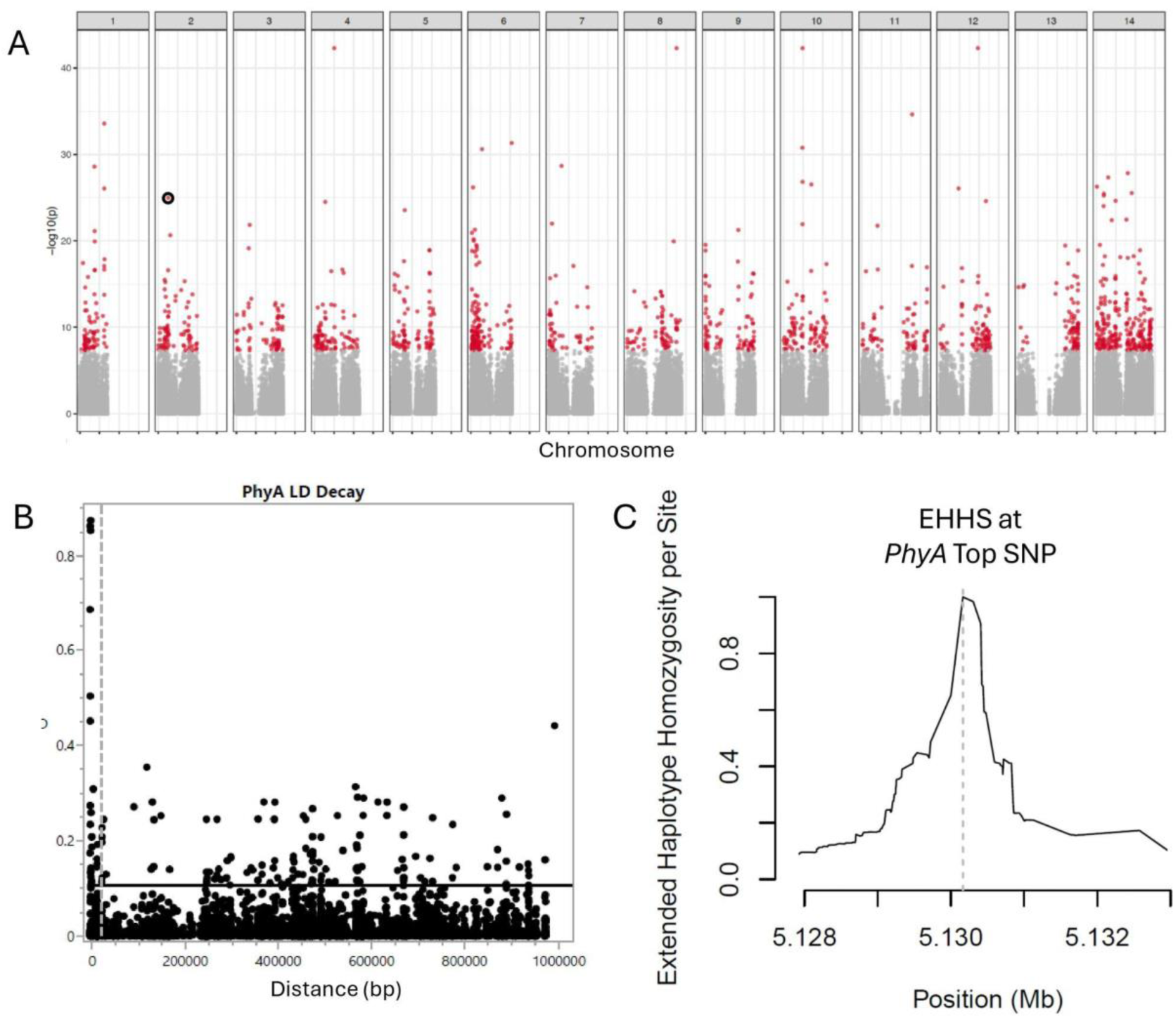
Signatures of selection on *PhyA*. (A) Whole genome manhattan plot of uncorrected –log10 Petiole Length LMM p_wld values. The most significant non-synonymous SNP [Chr_04:21534507] is circled in orange. The Bonferroni threshold is represented by the red line. (B) Full Range linkage disequilibrium (LD) decay plotted against the base-pair distance between the most significant non-synonymous SNP and nearby SNPs. Vertical dotted grey line denotes the full genome LD decay estimate. Horizontal black line denotes the threshold defining extended LD. (C) Full Range per site Extended Haplotype Homozygosity (EHH) around the most significant non-synonymous SNP.

**Figure S20:**
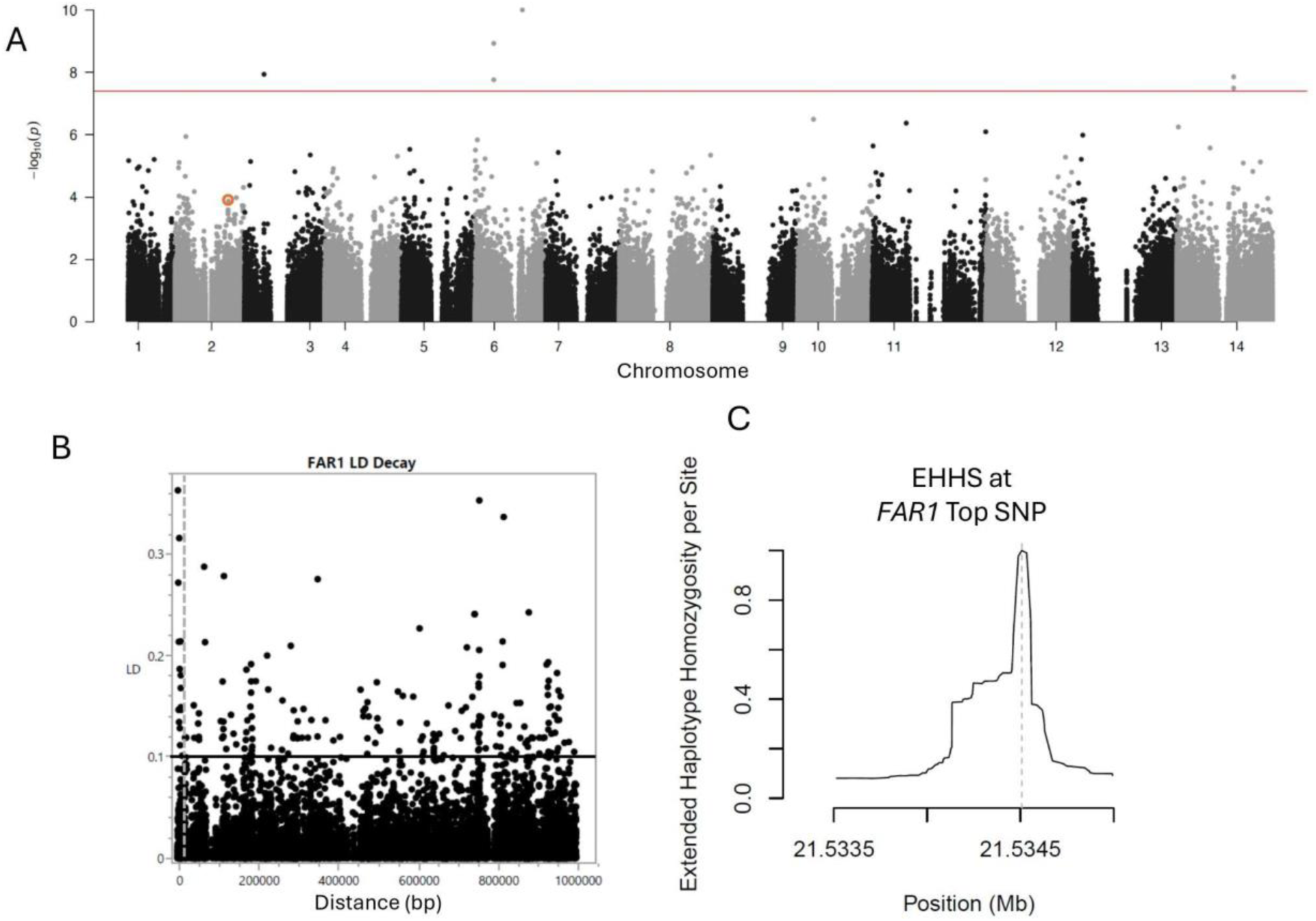
Signatures of selection on FAR1. (A) Whole genome manhattan plot of uncorrected –log10 Petiole Length LMM p_wld values. The most significant non-synonymous SNP [Chr_04:21534507] is circled in orange. The Bonferroni threshold is represented by the red line. (B) Full Range linkage disequilibrium (LD) decay plotted against the base-pair distance between the most significant non-synonymous SNP and nearby SNPs. Vertical dotted grey line denotes the full genome LD decay estimate. Horizontal black line denotes the threshold defining extended LD. (C) Full Range per site Extended Haplotype Homozygosity (EHH) around the most significant non-synonymous SNP.

**Figure S21:**
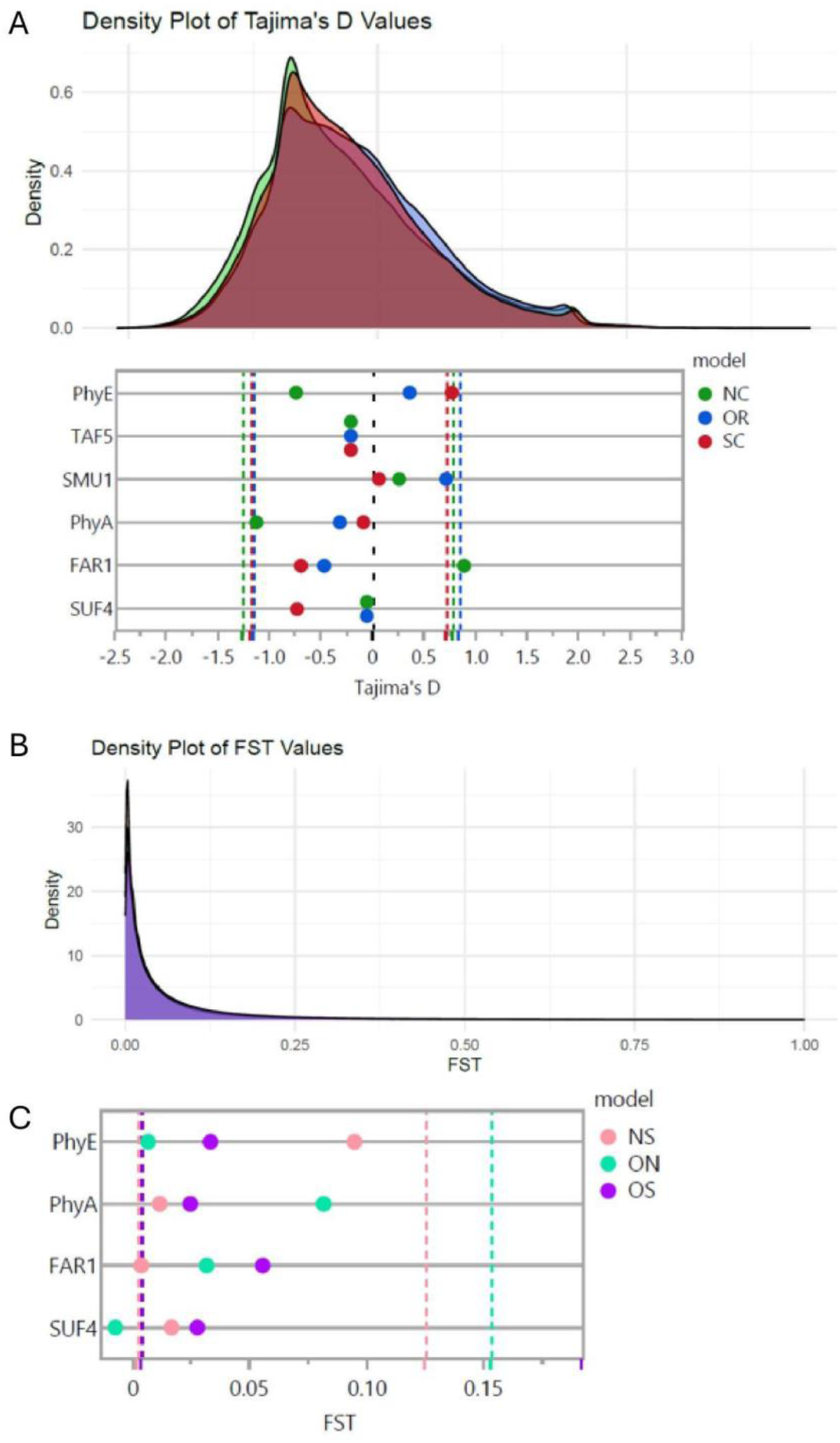
Signatures of selection for focal SNPs. (A) Overlapping density plot of GEA hit Tajima’s D values for each region. Underneath is the Tajima’s D values for each focal SNP colored by region. Vertical dotted lines represent the upper and lower 10% Tajima’s D values for all GEA hits by region. Upper density plot and lower dot plot are identically scaled, and values underneath directly represent their placement on the density plot above. (B) Overlapping density plot of GEA hit F_ST_ values for each region. Since F_ST_ is a pairwise metric, the comparisons between Oregon and Northern California are visualized as teal, Northern California and Southern California as coral, and Oregon and Southern California as purple. (C) F_ST_ values for each focal SNP colored by region using the same color scheme as B. Vertical dotted lines represent the upper and lower 10% F_ST_ values for all GEA hits by region. Upper density plot and lower dot plot are not identically scaled, and thus have their own x axis.

**Figure S22:**
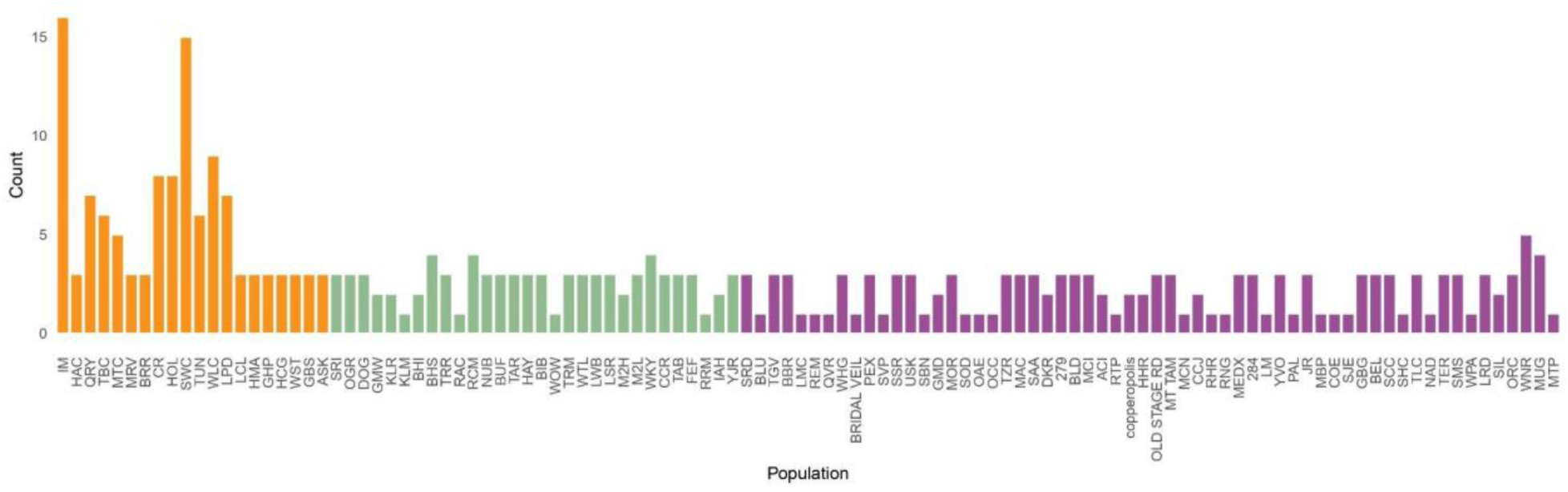
Distribution of individuals per population per region. Number of individuals per population colored by assigned region. Oregon is visualized in orange, Northern California is visualized in green, and Centraland Southern California is visualized in purple. The average number of individuals per population in Oregon is 5.85, in Northern California is 2.67 individuals, and Centraland Southern California is 2.17 individuals. Across the full range, the average number of individuals per population is 2.97 individuals.

**Figure S23:**
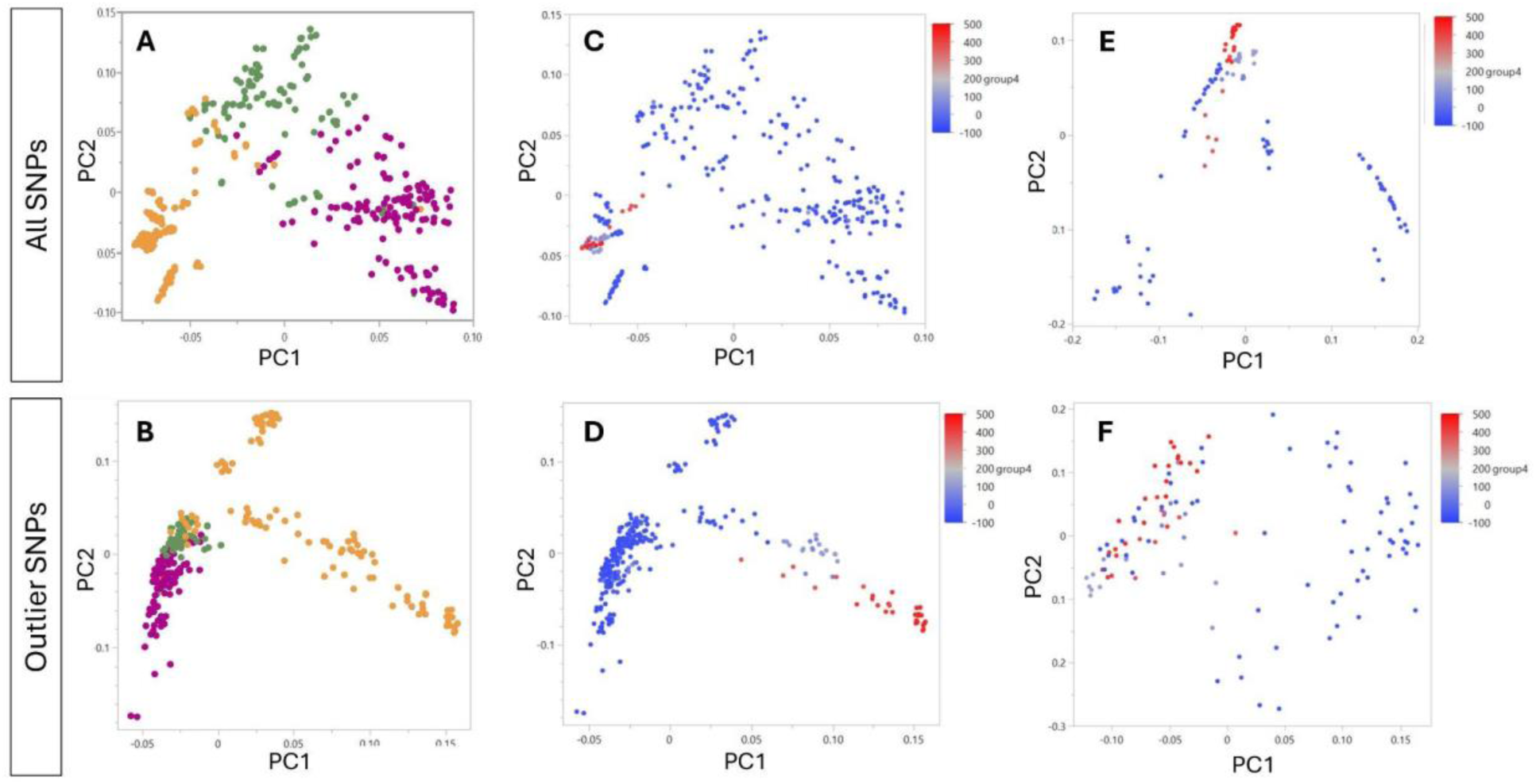
Full Range data skewed by Oregon, but population structure resolved in regional models. (A) Full range genetic variance (GenPC1, GenPC2) colored by region (Oregon = orange, Northern California = green, Central and Southern California = purple). (B) Full Range genetic variance for outlier SNPs only colored by region. (C) Full range genetic variance (GenPC1, GenPC2) colored by Group4 (“Snow”), the variable most responsible for the first axis of genetic variation across the full range. (D) Full range genetic variance for outlier SNPs only (GenPC1, GenPC2) colored by Group4 (“Snow”), the variable most responsible for the first axis of genetic variation across the full range. (E) Oregon genetic variance (GenPC1, GenPC2) colored by Group4 (“Snow”). (F) Oregon genetic variance for outlier SNPs only (GenPC1, GenPC2) colored by Group4 (“Snow”).

### Supplemental Tables

**Table S1: Sample history**. Sample provenance and year of collection. All sample reads were aligned and variants called as a combined dataset regardless of how they were obtained.

**Table S2: Range-wide environmental parameters**. BioClim variables were downloaded at a resolution of 30s from WorldClim (https://www.worldclim.org/). The 6 “group” variables consist of the PC1 value for each correlated variable grouping found in Table S3.

**Table S3: Final environmental variables for GEA analysis.** Final 17 Environmental Variables used for RDA and LFMM GEA analysis. All Variables were used for all regional models. Grouped Variables consist of the PC1 values for the listed variables combined, defined as variables with a within-cluster R^2^ > 0.7 as determined by JMP v 17pro variable clustering.

**Table S4: EnvPC and GenPC Individual Sample Values.** Individual sample values for the first three environmental PC values and the first two genetic PC values.

**Table S5: EnvPC x GenPC correlations**. Correlation between the first three environmental PC values and the first two genetic PC values for each regional model.

**Table S6: Correlations of GenPCX and EnvPCX per region**. R^2^ values of correlations between the first two genetic PC values and the first three environmental PC values by regional model.

**Table S7. Comparative landscape genomic modeling results evaluated via Multiple Matrix Regression with Randomization (MMRR) and Generalized Dissimilarity Modeling (GDM).** For the MMRR model, the coefficient β represents the standard linear regression matrix weights. For the non-linear GDM model, the metric M₂ represents the maximum I-spline height, indicating the total cumulative biological turnover (permutation variable importance) across the sampled environmental gradient or geographic distance matrix. Model performance is reported using R² (MMRR) and Percent Deviance Explained (GDM). Significance values (p) for both frameworks were derived using 500 permutations.

**Table S8: Amount of variance attributed to Environment increases across all regional models.** Variance partitioning of the amount of genetic variance attributable to each included metric using Full Range data (FR), the Oregon region (OR), the Northern California region (NC), and the Southern California region (SC). The full model uses the first two axes of genetic variation (GenPC1 and GenPC2), gps population coordinates, and the first three axes of environmental variance (EnvPC1, EnvPC2, and EnvPC3) as predictor variables in a partial RDA model. The pure environmental model uses the first three axes of environmental variance (EnvPC1, EnvPC2, and EnvPC3) as a predictor variable, and the model uses the first two axes of genetic variation (GenPC1 and GenPC2) and gps population coordinates as covariates. The pure population structure model uses the first two axes of genetic variation (GenPC1 and GenPC2) as a predictor variable, and the model uses the first three axes of environmental variance (EnvPC1, EnvPC2, and EnvPC3) and gps population coordinates as covariates. The pure geography model uses the gps population coordinates as a predictor variable, and the model uses the first three axes of environmental variance (EnvPC1, EnvPC2, and EnvPC3) and the first two axes of genetic variation (GenPC1 and GenPC2) as covariates. The proportion of explainable variance represents the amount of genetic variation that can be attributable to the predictor variables alone. Confounded variance includes variation due to mixtures of the included variables. The proportion of total variance also included genetic variance that cannot be explained by any part of the model inputs. Confounded variance includes variation due to mixtures of the included variables. Proportion of Explained Variance and Proportion of Total Variance are denoted as PEV and PTV, respectively.

**Table S9: Amount of variance uniquely attributed to BCM parameters.** Variance partitioning of the amount of genetic variance attributable to each included metric using Full Range data (FR). The full model uses the first two axes of genetic variation (GenPC1 and GenPC2), gps population coordinates, and the first three axes of environmental variance (EnvPC1, EnvPC2, and EnvPC3) as predictor variables in a partial RDA model. The pure environmental model uses the first three axes of environmental variance (EnvPC1, EnvPC2, and EnvPC3) as a predictor variable, and the model uses the first two axes of genetic variation (GenPC1 and GenPC2) and gps population coordinates as covariates. The pure population structure model uses the first two axes of genetic variation (GenPC1 and GenPC2) as a predictor variable, and the model uses the first three axes of environmental variance (EnvPC1, EnvPC2, and EnvPC3) and gps population coordinates as covariates. The pure geography model uses the gps population coordinates as a predictor variable, and the model uses the first three axes of environmental variance (EnvPC1, EnvPC2, and EnvPC3) and the first two axes of genetic variation (GenPC1 and GenPC2) as covariates. The proportion of explainable variance represents the amount of genetic variation that can be attributable to the predictor variables alone. Confounded variance includes variation due to mixtures of the included variables. The proportion of total variance also included genetic variance that cannot be explained by any part of the model inputs. Confounded variance includes variation due to mixtures of the included variables.

**Table S10: Environmental variables responsible for largest axis of genetic variation**. The original environmental variables most responsible for variation along the first two axes of variation and subsequently the first three RDA reduced dimension axes.

**Table S11: RDA outlier SNPs**. Combined RDA and WZA summary statistics for each outlier SNP with an RDA p value < 0.05 after Bonferroni correction and window WZA p value < 0.05. geneID and geneName annotations were derived from the *Mimulus guttatus* IM63 v3.1 refernce genome (Phytozome v13; https://phytozome-next.jgi.doe.gov/info/Mguttatusvar_IM62_v3_1). AtGeneName and AtGeneName annotations were derived from the *Arabidopsis thaliana* TAIR 10 refernce genome (Phytozome v13; https://phytozome-next.jgi.doe.gov/info/Athaliana_TAIR10). EFFECT and IMPACT values were determined usinf SnpSift and SnpEff. GO terms were additionally extracted using the *Mimulus guttatus* IM62 v3.1 reference genome.

**Table S12: GO term enrichment by method and model**. Gene Ontology terms were obtained from each gene using the *Arabidopsis thaliana* TAIR 10 reference genome (Phytozome v 13; https://phytozome-next.jgi.doe.gov/info/Athaliana_TAIR10). GO terms enrichment analysis using the Gene Ontology Resource (https://geneontology.org/) and Arabidopsis thaliana TAIR 10 refernce genome was then performed for all GO term categories for each method and model combination to select terms with a p value < 0.05 after FDR correction.

**Table S13: LFMM SNP hits**. Combined LFMM and WZA summary statistics for each outlier SNP with an LFMM adjusted p value < 0.05 after Bonferroni correction and window WZA p value < 0.05. geneID and geneName annotations were derived from the *Mimulus guttatus* IM63 v3.1 refernce genome (Phytozome v13; https://phytozome-next.jgi.doe.gov/info/Mguttatusvar_IM62_v3_1). AtGeneName and AtGeneName annotations were derived from the *Arabidopsis thaliana* TAIR 10 refernce genome (Phytozome v13; https://phytozome-next.jgi.doe.gov/info/Athaliana_TAIR10). EFFECT and IMPACT values were determined usinf SnpSift and SnpEff. GO terms were additionally extracted using the *Mimulus guttatus* IM62 v3.1 reference genome.

**Table S14: Overlapping GEA RDA and LFMM gene outliers.** Combined RDA and WZA summary statistics for each outlier SNP with an RDA p value < 0.05 after Bonferroni correction, LFMM adjusted p value < 0.05 after Bonferroni correction, and window WZA p value < 0.05. geneID and geneName annotations were derived from the *Mimulus guttatus* IM63 v3.1 refernce genome (Phytozome v13; https://phytozome-next.jgi.doe.gov/info/Mguttatusvar_IM62_v3_1). AtGeneName and AtGeneName annotations were derived from the *Arabidopsis thaliana* TAIR 10 refernce genome (Phytozome v13; https://phytozome-next.jgi.doe.gov/info/Athaliana_TAIR10). EFFECT and IMPACT values were determined usinf SnpSift and SnpEff. GO terms were additionally extracted using the *Mimulus guttatus* IM62 v3.1 reference genome.

**Table S15: Correlations between phenotypes and environmental variables.** Regression R^2^ values between all environmental variables and measured phenotypic traits from common garden.

**Table S16: GWA SNP hits.** Combined LMM and BSLMM summary statistics for each outlier SNP with an LMM p_wld > 0.05 after Bonferroni correction and BSLMM posterior probability (gamma) < 0.05. geneID and geneName annotations were derived from the *Mimulus guttatus* IM63 v3.1 reference genome (Phytozome v13; https://phytozome-next.jgi.doe.gov/info/Mguttatusvar_IM62_v3_1). AtGeneName and AtGeneName annotations were derived from the *Arabidopsis thaliana* TAIR 10 reference genome (Phytozome v13; https://phytozome-next.jgi.doe.gov/info/Athaliana_TAIR10). EFFECT and IMPACT values were determined using SnpSift and SnpEff. GO terms were additionally extracted using the *Mimulus guttatus* IM62 v3.1 reference genome.

**Table S17: Overlapping GEA LFMM and GWA hits.** Overlapping LFMM/WZA and LMM/BSLMM summary statistics for each outlier SNP with an LFMM adjusted pvalue < 0.05 after Bonferroni correction, WZA p value <0.05, LMM p_wld < 0.05 after Bonferroni correction, and BSLMM posterior probability (gamma) < 0.05. geneID and geneName annotations were derived from the *Mimulus guttatus* IM63 v3.1 refernce genome (Phytozome v13; https://phytozome-next.jgi.doe.gov/info/Mguttatusvar_IM62_v3_1). AtGeneName and AtGeneName annotations were derived from the *Arabidopsis thaliana* TAIR 10 refernce genome (Phytozome v13; https://phytozome-next.jgi.doe.gov/info/Athaliana_TAIR10). EFFECT and IMPACT values were determined usinf SnpSift and SnpEff. GO terms were additionally extracted using the *Mimulus guttatus* IM62 v3.1 reference genome.

